# Experimental and computational study on motor control and recovery after stroke: towards a constructive loop between experimental and virtual embodied neuroscience

**DOI:** 10.1101/2020.04.22.019661

**Authors:** Anna Letizia Allegra Mascaro, Egidio Falotico, Spase Petkoski, Maria Pasquini, Lorenzo Vannucci, Núria Tort-Colet, Emilia Conti, Francesco Resta, Cristina Spalletti, Shravan Tata Ramalingasetty, Axel von Arnim, Emanuele Formento, Emmanouil Angelidis, Camilla Hagen Blixhavn, Trygve Brauns Leergaard, Matteo Caleo, Alain Destexhe, Auke Ijspeert, Silvestro Micera, Cecilia Laschi, Viktor Jirsa, Marc-Oliver Gewaltig, Francesco S. Pavone

**Affiliations:** Neuroscience Institute, National Research Council, Pisa; European Laboratory for Non-Linear Spectroscopy, Sesto Fiorentino (Fi); The BioRobotics Institute, Scuola Superiore Sant’Anna, Pontedera (Pisa); Aix-Marseille Université, Inserm, INS UMR_S 1106, Marseille, France; Paris-Saclay University, Institute of Neuroscience, CNRS, Gif Sur Yvette, France; Department of Physics and Astronomy, University of Florence, Italy; Biorobotics Laboratory, École polytechnique fédérale de Lausanne (EPFL), Lausanne, Switzerland; Fortiss GmbH, Munich, Germany; Bertarelli Foundation Chair in Translational NeuroEngineering, Institute of Bioengineering, Swiss Federal Institute of Technology (EPFL), Lausanne, Switzerland; Chair of Robotics, Artificial Intelligence and Embedded Systems, Department of Informatics, Technical University of Munich, Germany; Institute of Basic Medical Sciences, University of Oslo, Norway; Department of Biomedical Sciences, University of Padua, Italy; Blue Brain Project (BBP), École polytechnique fédérale de Lausanne (EPFL), Genève, Switzerland

## Abstract

Being able to replicate real experiments with computational simulations is a unique opportunity to refine and validate models with experimental data and redesign the experiments based on simulations. However, since it is technically demanding to model all components of an experiment, traditional approaches to modeling reduce the experimental setups as much as possible. In this study, our goal is to replicate all the relevant features of an experiment on motor control and motor rehabilitation after stroke. To this aim, we propose an approach that allows continuous integration of new experimental data into a computational modeling framework. First, results show that we could reproduce experimental object displacement with high accuracy via the simulated embodiment in the virtual world by feeding a spinal cord model with experimental registration of the cortical activity. Second, by using computational models of multiple granularities, our preliminary results show the possibility of simulating several features of the brain after stroke, from the local alteration in neuronal activity to long-range connectivity remodeling. Finally, strategies are proposed to merge the two pipelines. We further suggest that additional models could be integrated into the framework thanks to the versatility of the proposed approach, thus allowing many researchers to achieve continuously improved experimental design.

## 1 INTRODUCTION

In nature, the activity of the brain of an individual interacting with the environment is conditioned by the response of the environment itself, in that the output of the brain is relevant only if it has the ability to impact the future and hence the input the brain receives. This “closed-loop” can be simulated in a virtual world, where simulated experiments reproduce actions (output from the brain) that have consequences (future input to the brain) (Zrenner et al., 2016). To the aim of reproducing *in silico* the complexity of real experiments, different levels of modeling shall be integrated. However, since modeling all components of an experiment is very difficult, traditional approaches of computational neuroscience reduce the experimental setups as much as possible. An “Embodied brain” (or “task dynamics”, see Zrenner et al. (2016)) approach could overcome these limits by associating the modelled brain activity with the generation of behavior within a virtual or real environment, i.e. an entailment between an output of the brain and a feedback signal into the brain (Tessadori et al., 2012; DeMarse et al., 2001; Reger et al., 2000). The experimenter can interfere with the flow of information between the neural system and environment on the one hand and the state and transition dynamics of the environment on the other. Closing the loop can be performed effectively by (i) validating the models on experimental data, and (ii) designing new experiments based on the hypotheses formulated by the simulations. On the example shown in Fig. 1, data on brain activity (be it, for instance, from electrophysiological recordings or imaging) and on the environment (e.g. by means of kinematic or dynamic measures) from the real experiment are used to feed the models of the *in silico* representation of the experiment. From a comparison of the real and model-based data, the features that are most important to replicate the real experiment are identified, and thus novel insights are generated (Fig. 1). To realize such a complex virtual system, many choices can be made, for instance on the brain model or spinal cord model that best represent the salient features of experimental measures to be replicated. The ideal framework shall comprise a library of tools to choose from, to reproduce a variety of experimental paradigms in the virtual environment. By briefly introducing the state of the art in brain and spinal cord modeling, we will discuss few classes of models to pick from an ideal library.

**Figure 1.**
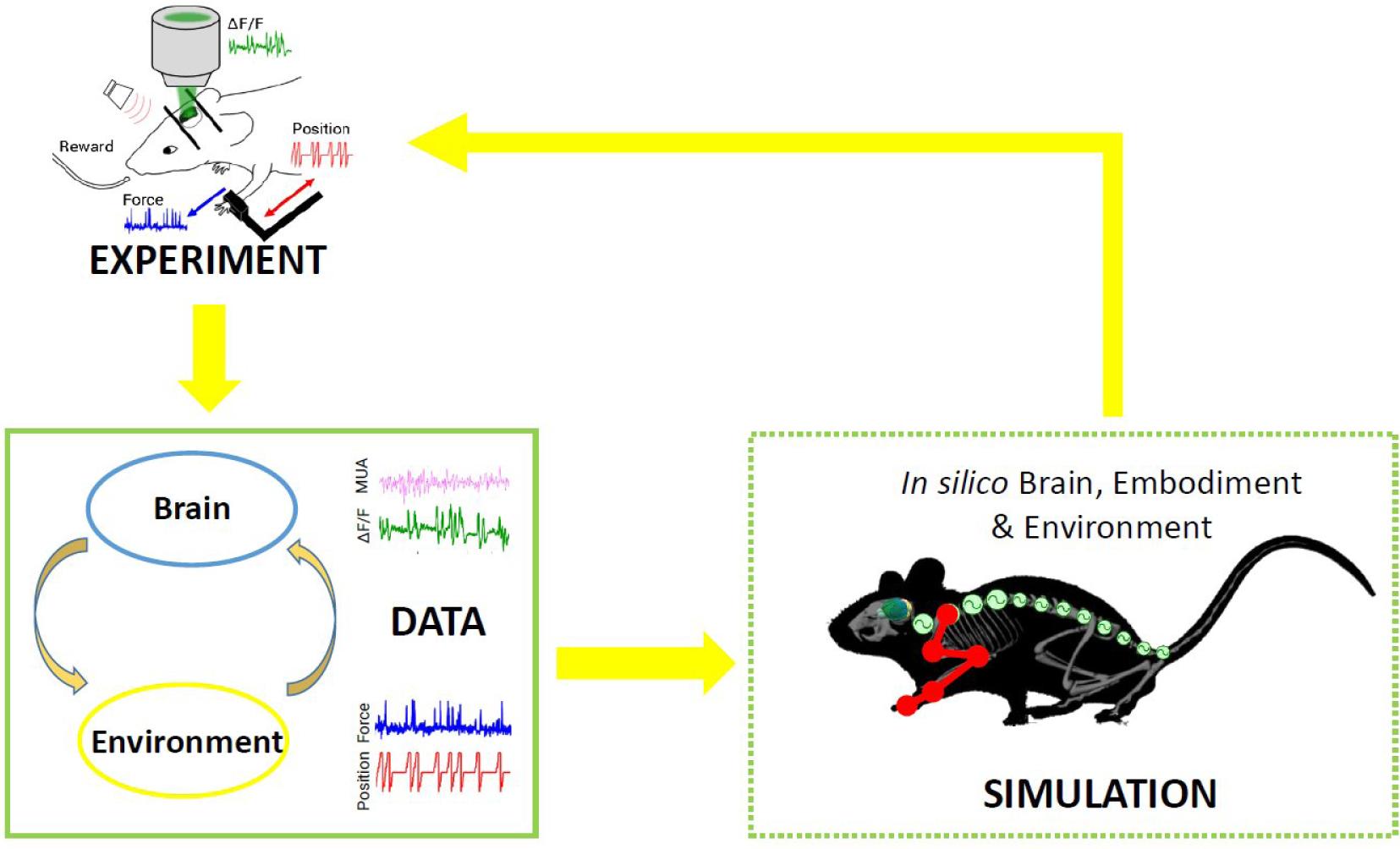
Scheme of the proposed *Embodied brain* framework. The picture suggests a closed-loop workflow linking real and simulated experiment. The different types of data obtained from the experiments, from brain activity to dynamic and kinematics of goal-directed movement, are used to feed the whole brain and spinal cord model, in addition to the virtual mouse and environment. The loop is closed by validation of *in silico* results on real data. Eventually, the simulated experiment raises novel hypotheses, to be validated on new real experiments.

### 1.1 State of the art

#### 1.1.1 Local cortical network modeling

Biologically detailed models of a single neuron such as the Hodgkin and Huxley model (Hodgkin and Huxley, 1952) take into account the activity of the ion channels in the cell membrane that lead to changes in the membrane potential, eventually causing the neuron to spike. However, simpler but still biologically realistic models of the single cell are preferable when interested in modeling the dynamics of a larger number of cells. A good candidate is the adaptive exponential integrate and fire (adex) neuron model (Brette and Gerstner, 2005), which has been shown to reproduce the intrinsic neuronal properties of a number of cell types, including those with spike frequency adaptation (Destexhe, 2009). Interestingly, adex neurons network models with different adaptation levels can reproduce the dynamical properties of distinctive brain states such as wakefulness, sleep or anesthesia (Zerlaut and Destexhe, 2017; Nghiem et al., 2020). This property makes adex neuron networks suitable to model the dynamics of the emerging activity in a local network of neurons after injury, given that, after a stroke, the dynamics of the local network switches to a slow oscillatory rhythm which resembles that of sleep or anesthesia Butz et al. (2004). Moreover, alterations in low-frequency cortical activity in the peri-infact cortex after stroke are known to correlate with motor recovery (e.g. Yilmaz et al. (2015); Ramanathan et al. (2018))

#### 1.1.2 Brain Network Modeling

Efforts have been made to reconstruct single brain regions with as many details as possible, (Markram et al., 2015), or to build detailed networks of multi-compartment oscillators (Izhikevich and Edelman, 2008). Contrary to the detailed models, top-down modelling seeks to elucidate whole-brain network mechanisms, which may underpin a variety of apparently diverse neurophysiological phenomena. Neural masses formalisms have been used over many years to develop macroscopic models that capture the collective dynamics of large neural assemblies (Deco et al., 2008; Sanz-Leon et al., 2015). In this case the activity of a macroscopic brain region is often directly derived from populations of spiking neurons as a mean-field using concepts from statistical physics, e.g. (Stefanescu and Jirsa, 2008; Wong and Wang, 2006; Zerlaut et al., 2017). In other cases the statistics of the macroscopic brain activity is derived more phenomenologically while still conserving some basic physiological principles such as division on excitatory and inhibitory neurons, e.g. the seminal Wilson Cowan model (Wilson and Cowan, 1972). The third subclass of neural masses contains purely phenomenologically derived computational models that aim to reproduce certain dynamical properties of the macroscopic neuronal activity, such as e.g. seizure dynamics (Jirsa et al., 2014; Saggio et al., 2017), whilst different realizations of damped or self-sustained oscillators are often used to model the coherent fluctuations of resting-state activity (Cabral et al., 2011; Deco et al., 2016). Depending on the working point of the system, the macroscopic dynamics can be described not only by physiologically derived mean-fields, but also by phenomenological models in their canonical form (Deco et al., 2009, 2011; Izhikevich, 1998). Hence, phase oscillators (Kuramoto, 1984) are often chosen to model and study the coactivation patterns in the brain, as some kind of a minimal model explanation (Batterman and Rice, 2014) for the synchronized behaviour over a network (Pikovsky et al., 2001; Breakspear et al., 2010).

Connecting the neural masses in large-scale brain network models (BNM) became possible with the progress of non-invasive structural brain imaging (Johansen-Berg and Rushworth, 2009). This allowed extraction of biologically realistic brain connectivity, the so-called connectome, which shapes the local neuronal activity to the emergent network dynamics (Honey et al., 2007; Ghosh et al., 2008; Deco et al., 2009; Sanz-Leon et al., 2015; Petkoski et al., 2018; Petkoski and Jirsa, 2019).

The large-scale BNM have been used to interpret healthy (Cabral et al., 2011; Deco et al., 2016) or pathological (Nakagawa et al., 2013; Saenger et al., 2017; Zimmermann et al., 2016) brain activity. This is often reflected in the coherence between brain rhythms (Lachaux et al., 1999) that also describes the functional connectivity (FC) of the brain as an important marker of its spatio-temporal organization (Deco et al., 2009, 2011; Ghosh et al., 2008; Deco and Jirsa, 2012; Petkoski et al., 2018).

The Virtual Brain (TVB) (Sanz Leon et al., 2013; Sanz-Leon et al., 2015) is a commonly used neuroinformatics platform for full brain simulations. It supports a systematic exploration of the underlying components of a large-scale BNM: the structural connectivity (SC) and the local dynamics that depend on the neurophysiological mechanisms or phenomena being studied. In this way, the BNM allows to describe structural changes (through connectivity variation including stroke, motor learning and recovery) and subsequent functional consequences accessible to modeling and empirical data collection on the meso, macro and behavioral level. The modelling with TVB thus represents a useful paradigm for multi-scale integration. TVB has been already utilized in modeling functional mechanism of recovery after stroke in humans (Falcon et al., 2015, 2016), identifying that the post-stroke brain favors excitation-over-inhibition and local-over-global dynamics. For studying the changes in synchronization, as we intend to do, TVB offers a range of oscillatory models for the neural activity. One of these is the Kuramoto model (KM), which captures the emergent behaviour of a large class of oscillators that are near an Andronov-Hopf bifurcation (Kuramoto, 1984), including some population rate models Ton et al. (2014). This makes the KM well suited for assessing how the connectome governs the ynchronization between distant brain regions (Breakspear et al., 2010; Cabral et al., 2011, 2012; Ponce-Alvarez et al., 2015; Petkoski et al., 2018).

#### 1.1.3 Spinal cord modeling

The brain controls its body through neural signals originating from the brain and processed by the spinal cord to control muscle activation in order to perform a large variety of behaviors. Several biologically realistic functional models of the spinal cord have been developed and tested in closed loop simulations with musculoskeletal embodiments. Stienen and colleagues (Stienen et al., 2007) developed a fairly complete model that includes Ia, Ib and II sensory afferents, both monosynaptic and polysynaptic reflexes as well as Renshaw cells, improving a previous work by Bashor (Bashor, 1998). The model was tested with a musculoskeletal model consisting of a generic antagonistic couple of muscles, thus lacking a realistic validation scenario. Cisi and Kohn developed a web-based framework for the simulation of generic spinal cord circuits with associated muscles, that aims at replicating realistic experimental conditions (i.e. electrical stimulation) (Cisi and Kohn, 2008). Sreenivasa and colleagues developed a specific neuro-musculoskeletal system, upper limb with biceps and triceps, and validated it against human recordings (Sreenivasa et al., 2016). In (Moraud et al., 2016), a simple spinal cord model of the rat, lacking any descending stimuli, was developed in order to study how such circuitry can correct the gait after a spinal cord injury and embedded in a closed loop simulation with biomechanical hindlimbs. All of the mentioned works were tested primary for the generation of reflex motions, and not as intermediate levels of more complex controllers such as ones capable of generating voluntary movements.

### 1.2 Aim of the work

We propose a framework (“*Embodied brain closed loop*”) endowed with a library of modeling tools that will eventually allow to realize entirely virtual experiments. We focused on an experiment on motor control and motor recovery after stroke described in Spalletti et al. (2017) and Allegra Mascaro et al. (2019), whose simulation requires two main tiles. The first is the realization of voluntary movements in a virtual milieu. This piece requires monitoring and modeling of many components of movement control, from brain activity to body kinematics and displacement of virtual objects. The second is the simulation of brain injury. This includes modeling of acute consequences but also of neuronal plasticity after brain damage, either spontaneous or supported by treatment. Both local and long-range modulation of neuronal activity should be accounted to simulate the brain after stroke, since local alteration of neuronal activity in the peri-infarct area is known to be associated to remodeling of long-range functional and structural connectivity (several comprehensive reviews have summarized this research, e.g. Carmichael et al. (2017)). To build those tiles, we developed two pipelines that target, on one side, the physiological execution of movements and, on the other, pathological alterations and plasticity (Fig. 2). The first (“Movement-driven models” pipeline) aims at reproducing in a virtual environment how a goal-directed movement is performed and represented in the healthy brain. Data recorded on healthy mice are used as an input to the spinal cord model, attached to the muscles of the simulated embodiment (see Fig. 2, red box). The goal of the second pipeline (“Stroke models”) is to reproduce both local and long-range consequences of stroke. We developed a spiking neurons model that could simulate the local brain dynamics, and in particular the abnormal oscillatory activity taking place in the peri-infarct cortex (see Fig. 2, lower line in the green box). Also, we show how the simulation of brain activity by neural mass models allows replicating the evolution of functional connectivity in mouse brain after a stroke and under rehabilitation (see Fig. 2, upper line in the green box).

**Figure 2.**
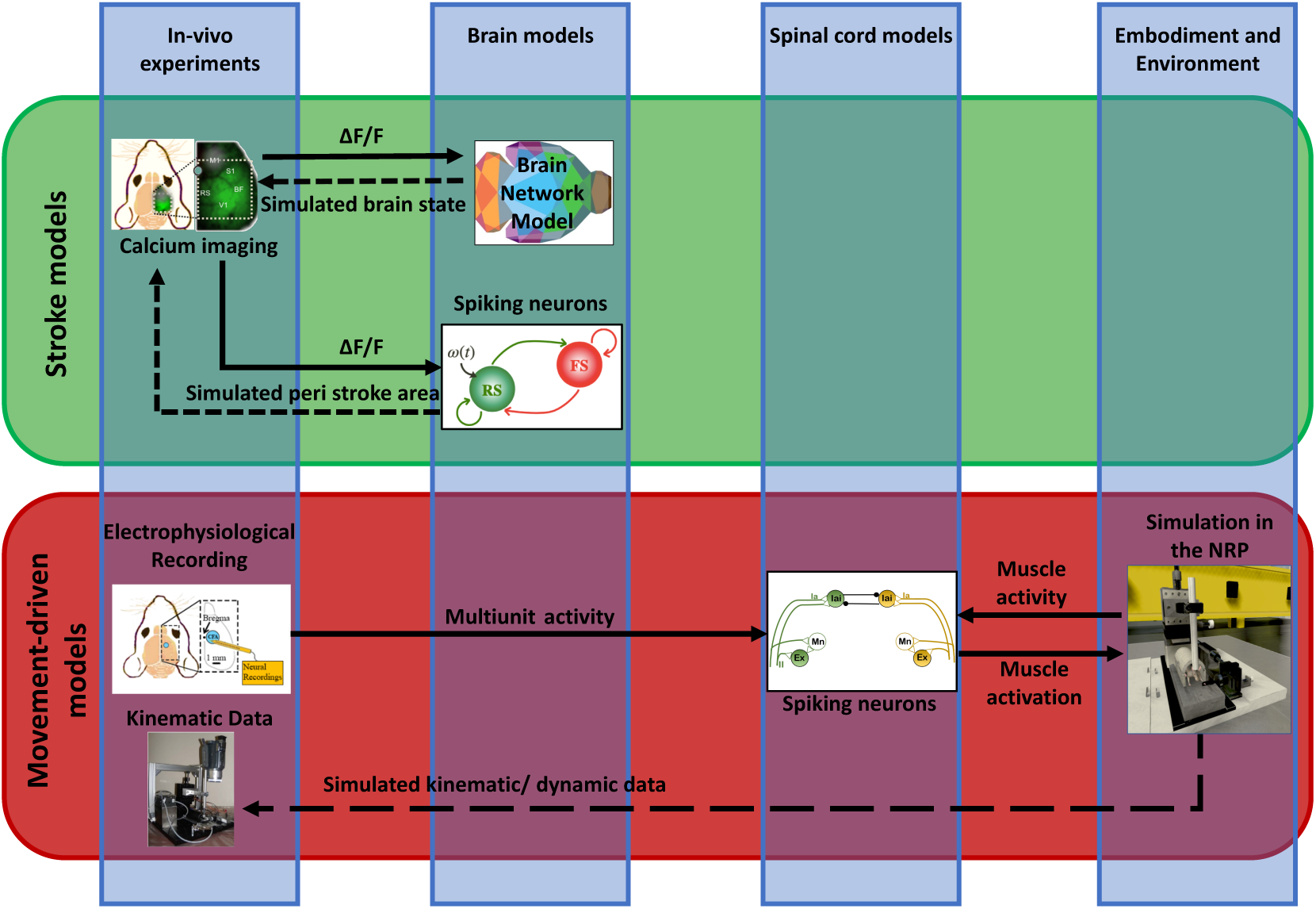
Scheme of data and simulations. The scheme depicts the approach to build the *Embodied brain* framework from data to models and back. The workflow from data to models and simulation in the *Embodied brain closed loops* is shown. The upper, green box shows the *Stroke models* closed loop, the lower, red box shows the *Movement-driven models* closed loop. Colored images represent experiment data, brain and spinal cord models, and simulation of the environment (from left to right). Connections between the modeling components are presented as arrows: solid lines represent the output provided to other blocks; dashed lines indicate the output data of the models that are used for comparison with real data for validation.

## 2 METHODS

Cortical recordings and behavioral data from the experiments described in this section are used to build and validate the brain models and the output in the virtual environment.

### 2.1 *in vivo* experiments

On the experimental side, we performed electrophysiological recordings (Fig. 3, panel A) and wide-field calcium imaging (Fig. 3, panel B) in awake mice performing active forelimb retraction on a robotic device (M-Platform). These experiments allowed gathering simultaneous information on the neuronal activity, force applied during active forelimb retraction and position of the forelimb, as displayed in the lower panels of figure 3. The electrophysiological data and the recordings of limb position were used to feed the spinal cord model, as described in section 4.1. The features of the wide-field calcium data recordings were used to build the spiking neurons brain model and to validate the BNM, Section 3.3. All the procedures were in accordance with the Italian Ministry of Health for care and maintenance of laboratory animals (law 116/92) and in compliance with the European Communities Council Directive n. 2010/63/EU, under authorizations n. 183/2016-PR (imaging experiments) and n. 753/2015-PR (electrophysiology experiments).

**Figure 3.**
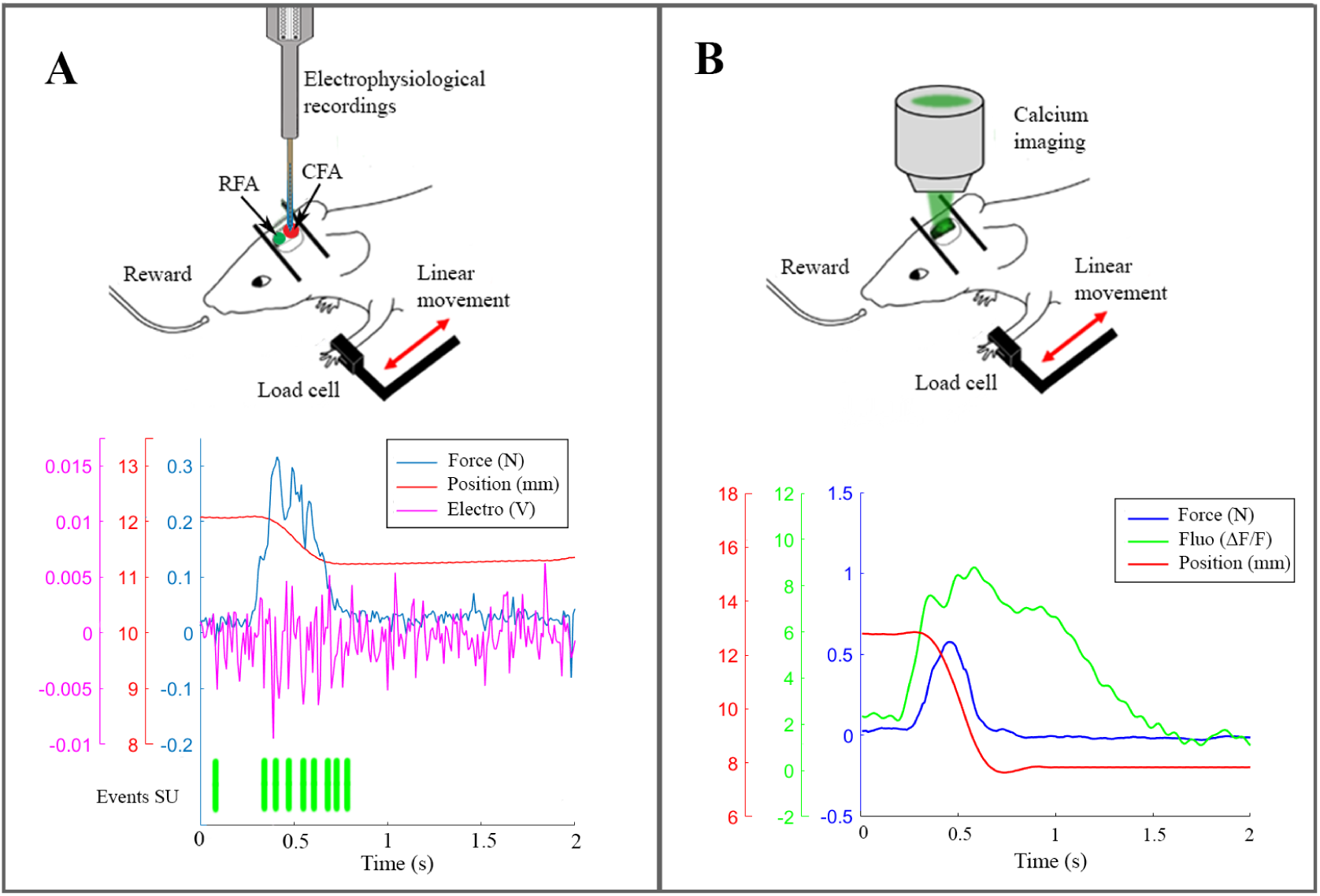
The real robotic platform. (A) On the top a schematic representation of the experiment during electrophysiological recording in CFA. At the bottom the synchronized data: force peak (blue), movement of the slide (red), high frequency electrophysiological signal of a single channel (magenta) and the timestamp of a selected single unit. (B) On the top a scheme of the experiment with the setup to record calcium activity. At the bottom the recorded data after synchronization: force peak (blue), movement of the slide (red) and calcium response (green).

#### 2.1.1 Robotic training on the M-Platform

The M-Platform is a robotic device designed to train mice to perform active forelimb retraction (Pasquini et al., 2018). Briefly, the main component of the device is a linear actuator that moves a linear slide where a custom handle is screwed. Moreover, the platform is provided with a system to control the friction on the slide and a pump for the reward. During the experiments, while the mouse has its left paw connected to the slide, first the linear actuator extends the forelimb then the animal has to perform an active pulling movement to come back to the starting point and to receive a reward. Force signal and position of the forelimb are recorded respectively by a load cell and a webcam.

In Section 2.1.3 and 2.1.4 we describe two different experiments with the robotic device. In the first one, the M-Platform is embedded with Omniplex D System (Plexon, Usa) to obtain *in vivo* electrophysiological recording during the task. In the second one, the kinetic and kinematic parameters are synchronized with wide-field calcium imaging recordings (Fig. 3).

#### 2.1.2 Photothrombotic Stroke

To induce focal stroke in the right hemisphere, mice were injected with Rose Bengal (0.2 ml, 10 mg/ml solution in Phosphate Buffer Saline). Five minutes after intraperitoneal injection, a white light from an LED lamp was focused with a 20X objective and used to illuminate the primary motor cortex (0.5 mm anterior and 1.75 mm lateral from Bregma) for 15 min.

#### 2.1.3 Electrophysiological recordings on the M-Platform

Two healthy mice were used for the experiments. Animals were housed on a 12h/12h light/dark cycle. Mice were water deprived overnight before training on the platform; daily liquid supplement was given after the test. Food was available ad libitum. To have access to the motor cortex, a craniotomy was performed 3 days before the training to expose the Caudal Forelimb Area (CFA) of the right hemisphere.The craniotomy was filled with agarose and silicon (Kwik cast sealant, WPI) and could be opened and closed several times for acute recordings.”

Mice were gradually acclimated to the platform. Then they performed the task for two days, fifteen trials each day. During the pulling experiment, mice were head fixed to the platform with their left wrist constrained to the slide. The friction on the slide was set at 0.3 N. The force signal was acquired by a load cell (Futek LSB200, CA, USA) along the direction of the movement at 100 Hz, at the same time a webcam recorded the position of the slide at 25 Hz and the multi-unit activity was recorded by Omniplex D System (Plexon, USA) with a frequency of 40 kHz thanks to a 16 channels linear probe (1 *M* Ω,ATLAS, Belgium) inserted into the CFA at 850 *µm* of depth. (Fig. 3A).

#### 2.1.4 Wide-field calcium imaging of cortical activity during training on the M-Platform

The mouse was housed in clear plastic cage under a 12 h light/dark cycle and was given ad libitum access to water and food. We used the following mouse line from Jackson Laboratories (Bar Harbor, Maine USA): C57BL/6J-Tg(Thy1GCaMP6f)GP5.17Dkim/J (referred to as GCaMP6f mice). In this mouse model, the fluorescence indicator GCaMP6f is mainly expressed in excitatory neurons H. Dana (2014). GCaMP6f protein is ultra-sensitive to calcium ions concentration T.W. Chen and Kim (2013); H. Dana (2014) whose increase is associated with neuronal firing activity Grienberger and Konnerth (2012); R. Yasuda and Svoboda (2004).

For wide-field fluorescence imaging of GCaMP6f fluorescence, we used a custom made microscope described in Conti et al. (2019). Briefly, the system is composed by a 505 nm LED (M505L3 Thorlabs, New Jersey, United States) light deflected by a dichroic filter (DC FF 495-DI02 Semrock, Rochester, New York USA) on the objective (2.5x EC Plan Neofluar, NA 0.085, Carl Zeiss Microscopy, Oberkochen, Germany). The fluorescence signal is selected by a band pass filter (525/50 Semrock, Rochester, New York USA) and collected on the sensor of a high-speed complementary metal-oxide semiconductor (CMOS) camera (Orca Flash 4.0 Hamamatsu Photonics, NJ, USA).

The experiment starts with a mouse being trained and recorded for one week (five days) on the M-platform (“healthy” condition, see Fig. 3). The focal stroke is then induced at the beginning of the second week by phototrombosis on the right primary motor cortex (rM1). Starting 26 days after stroke, the mouse performance and spontaneous motor remapping was evaluated on the M-Platform for 5 days a week along 4 more weeks. The results from the first week one month after the injury is the so-called “stroke” condition, while the results during the last week, when the animal recovers the motor function is referred to as “rehab”.

Each day, the beginning of the wide-field imaging session was triggered by the start of the training session on the M-Platform. To detect the movement of the wrist of the animal in the low-light condition of the experiment, an infrared (IR) emitter was placed on the linear slide, and rigidly connected to the load cell and thus to the animal’s wrist. Slide displacement was recorded by an IR camera (EXIS WEBCAM #17003, Trust) that was placed perpendicular to the antero-posterior axis of the movement. Position and speed signals were subsequently extracted from the video recordings and synchronized with the force signals recorded by the load cell (sampling frequency = 100 Hz) and with the fluorescence signal recorded by the CMOS sensor (Fig. 3B).

### 2.2 Data analysis

#### 2.2.1 Spikes and force analysis

Data were analyzed offline using custom routines in Matlab (MathWorks). First, the position signal was extracted by the video using a white squared marker on the slide as reference. The recording frequency of the video was 25 Hz. After applying an antialiasing FIR lowpass filter, a uniform linear resample of the movement of the slide was performed, in order to synchronize the position signal with the force data, recorded at 100 Hz. To identify the timing of the voluntary activity of the animal, a threshold method was used to detect force peaks during the pulling phase of the task. For the following analysis, we picked out peaks that produced a displacement of the slide, in addition to crossing the threshold; and we calculate the onset of these peaks as the minimum of the force derivative just before the respective peak (Spalletti et al., 2014). The electrophysiological signal, recorded at 40 kHz as sampling rate, was analysed by Offline Sorter (Plexon, Dallas, TX). First, for each channel of the probe, we sorted waveform that crossed a detection threshold of the mean *±*3 standard deviations. Then, detected spikes were clustered using an automatic process based on principal component analysis. Starting from these clusters, a manual sorting was executed to isolate all single units which could be identified in the recorded multi-units signal. The time stamp of each unit was synchronized with the data of the robot. To evaluate the temporal behavior-related spike activity, the peri-stimulus time histograms (PSTHs, NeuroExplorer, Plexon) was generated with bins of 20 ms in an interval of 1 s around the onset of force peaks. In addition, the resting activity of each unit was evaluated selecting intervals of at least 0.6 s with no force peaks and calculating the average of the number of spikes in bins of 20 ms. Finally, the PSTHs was used to evaluated when a single neuron was active, that is when the number of spikes for bin cross the threshold, calculated as the mean *±*2 standard deviations of the number of spikes for bin during the respective resting activity.

#### 2.2.2 phase coherence and Funtional connectivity

Functional connectivity (FC) among cortical regions was inferred from phase coherence of activity measurements, and used to determine changes in brain activity in “stroke” and “rehab” condition, as compared to the healthy mice. These inferred activity changes were used to parameterize simulations of the BNM built over the Allen Brain Atlas mouse connectivity data (http://connectivity.brain-map.org/; (Oh et al., 2014), below referred to as the Allen Mouse Brain Atlas - AMBA), incorporated in the extended virtual mouse brain (Melozzi et al., 2017). In each animal, the camera field-of-view used for activity measurements was placed in a standard position using the sagittal suture and its intersection with the coronal suture of the skull (bregma) as anatomical landmarks. To spatially correlate our activity measures with the structural connectivity data (Oh et al., 2014), the camera field-of-view (Fig. 3, panel B) was spatially translated to the Allen Common Coordinate Framework (CCF, v3, 2015; (Oh et al., 2014)). Since the CCF lacks stereotactic skull landmarks, these were introduced by spatially co-registering all diagrams from a standard stereotaxic mouse brain atlas (Franklin et al., 2008) to the CCF coordinate space with affine transformations defined using the QuickNii tool (Puchades et al., 2019). Using bregma and the sagittal suture as a reference, the four corners of the downsampled 128×128 pixels field-of-view of the recorded images were positioned in CCF, taking the 5 degree lateral tilt of the camera view into account. Delineations of layer IV cortical regions were then projected onto the camera field-of-view, and used as a custom atlas reference for all activity maps.

The spectral content of the signals is analyzed to identify the frequency band which captures the spontaneous brain activity that occurs simultaneously with the motor-evoked events. Even though the slowest dynamics *<* 0.5*Hz*, which has the highest power, is often a marker of the resting state (Wright et al., 2017), in this experiment it also contains the propagation of waves generated during the limb movements on the platform. The mechanisms behind stimulation propagation (Spiegler et al., 2016) are different from the spontaneous oscillations at rest (Deco and Jirsa, 2012) that we try to study and model here, and hence the lowest frequencies are excluded from the analysis. In addition, the mice heart rate is between 6 − 8*Hz*, whilst the activity above 10*Hz* is too close to the Nyquist frequency of 12.5*Hz*, defined as half of the sampling rate of the recordings. As a consequence these bands are generally avoided in the analysis of calcium signals, which is consequently often centered at the *δ* band between around 1 Hz and 5 Hz, (Vanni et al., 2017; Wright et al., 2017).

The FC is characterized with the phase coherence of the analytical phases of the band-passed time-series obtained using the Hilbert transform (Pikovsky et al., 2001). For this we employ phase locking values (PLV) (Lachaux et al., 1999) that are a statistical measure for similarity between the phases of two signals, hence defined as

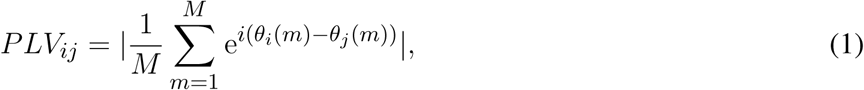

where the phase difference *θ*_*i*_(*m*) − *θ*_*j*_(*m*) between the regions *i* and *j* is calculated at times *m* = 1 *… M*. The same procedure is also applied to surrogate time-series to find the level of statistically significant phase coherence (Lancaster et al., 2018).

## 3 MODELS

### 3.1 Spinal cord model

To develop the final model, an incremental approach was followed, starting from a circuit for a single *muscle*, adding inhibitory connections between *antagonistic* pairs and finally interneurons to modulate descending stimuli (Fig. 4). For a single muscle, a network with muscle spindles providing Ia and II afferent fibers activity, a pool of *α*-motoneurons and excitatory II-interneurons was considered (Stienen et al., 2007; Moraud et al., 2016). Ia afferents directly provide excitatory inputs to the *α*-motoneurons (monosynaptic stretch reflex mechanism), while the II afferents output is mediated by a set of interneurons before reaching the *α*-motoneurons, creating a disynaptic reflex. The muscle spindles are implemented using the model from (Vannucci et al., 2017). All other neurons are modeled as leaky integrate and fire neurons. The number of neurons in the spinal cord populations, as well as parameters for the synaptic connections are taken from (Moraud et al., 2016), with the exception of the synaptic weights of the monosynaptic connections, which have been significantly lowered (see Supplementary Material). The parameters from the muscle spindle models are taken from (Mileusnic et al., 2006), which are tuned on neurophisiological recordings of lower mammals. Distribution of parameters for the *α*-motoneurons that influence the recruitment order and fiber strength (membrane capacitance, membrane time constant, maximum twitch force, time to peak force) are taken from (Sreenivasa et al., 2016):

**Figure 4.**
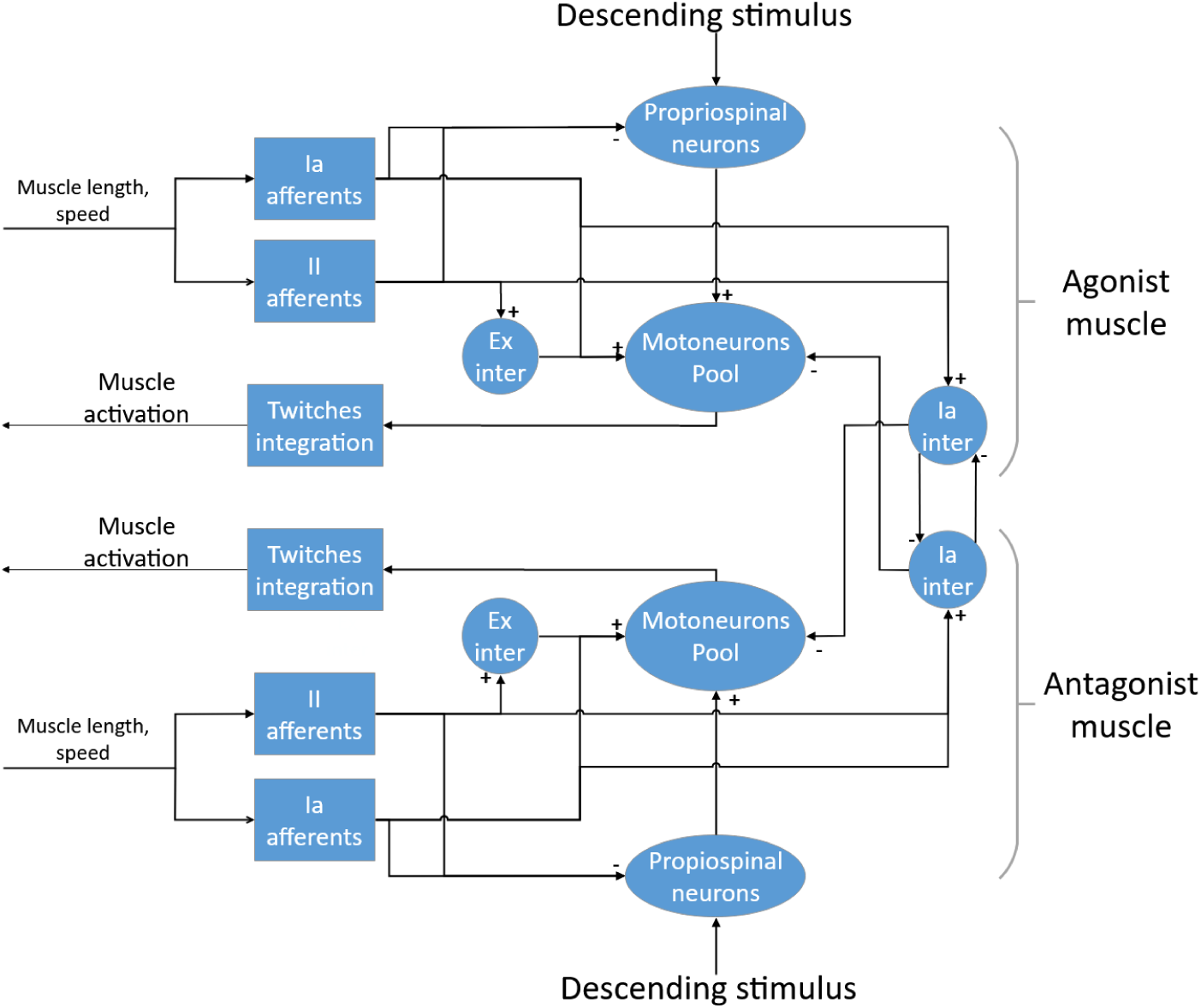
The spinal cord model for a pair of antagonistic muscles.

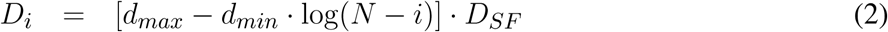

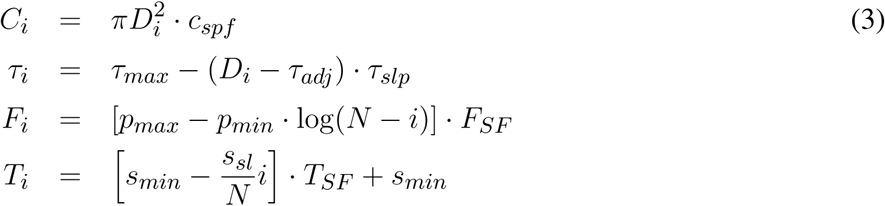

where *i* is the index of the *α*-motoneuron in the pool, *N* is the size of the pool and the others are free parameters that can be adjusted for every muscle. In this work, the value of these parameters has not been changed from (Sreenivasa et al., 2016).

In order to compute the actual muscle activation from the motoneurons activity, a special spike integration unit that sums the fibers twitches was implemented. The spikes were integrated using the discrete time equations of (Cisi and Kohn, 2008) with a non-linear scaling factor from (Fuglevand et al., 1993) that prevents the activation to grow indefinitely:

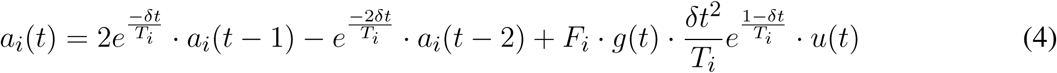

where *δt* is the integration time, and *u*(*t*) and *g*(*t*) are the spike function and the non-linear scaling, defined as:

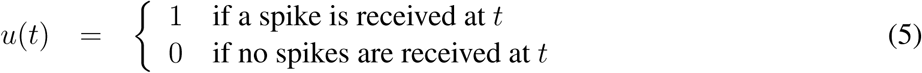

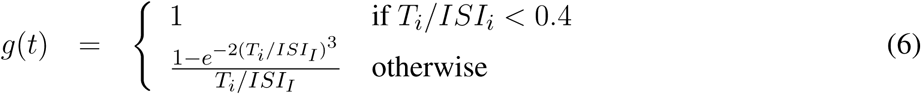

where *ISI*_*i*_ is the observed inter-spike interval of *α*-motoneuron *i*. Moreover, the activation can be scaled between 0 and 1 by dividing by the maximum theoretical value:

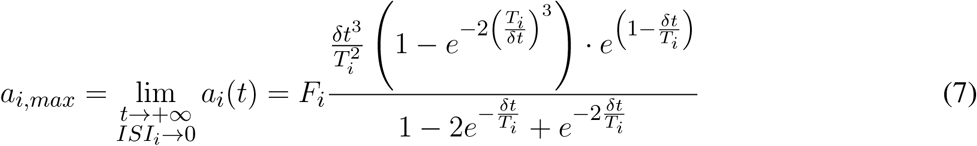

Therefore, the output of the twitch integration module is an activation value in [0; 1] that is suitable for the muscle model present on the mouse virtual embodiment. The effect of this integration is that at low frequencies the individual twitches can still be seen, while at higher stimulation the twitches fuse into a tetanic contraction. Moreover, thanks to the non-linear scaling, the activation reaches a maximum value and higher stimulation frequencies do not produce any effect, in accordance with the contractile properties of real fibers.

In order to implement the polysynaptic inhibition reflex between antagonistic muscles, two populations of Ia-interneurons were added to the network. Those receive inputs from all Ia afferents of a synergistic muscle and provide inhibition to the *α*-motoneurons of the corresponding antagonistic muscle. Moreover, as the activation of a muscle should provoke an inhibition of its antagonist (Pierrot-Deseilligny and Burke, 2005), the Ia-interneurons also receive low-gain positive inputs from the corresponding descending pathways. Again, the number of neurons in these population and their parameters have been taken from Moraud et al. (2016). Finally, as there is lack of evidence for a direct connection between cortical neurons and motoneurons in the spinal cord of rodents (Yang and Lemon, 2003), an intermediate population of neurons mediating descending signals was added to the circuitry. This population aims at modelling propriospinal neurons, which provide an inhibitory action on the signals coming from the corticospinal tract (Alstermark, 1992). In general, the inhibition is generated from different peripheral afferents, but we included only afferents from muscle spindles as these are the only present in the model. As there is no definitive experimental evidence on the size of the population of propriospinal neurons and its parameters, we set the values of these to those of the populations of Ia-interneurons. Conversely, the synaptic weights and the number of connections between the descending inputs and the propriospinal interneurons were empirically tuned starting from experimental data.

### 3.2 Simulation tools and physical models

This section describes simulation tools that were used to synchronize neural and physical simulations and the physical simulations models that have been developed and used. These tools and model were used in the context of the *Movement-driven models* pipeline.

#### 3.2.1 Embodied mouse in the Neurorobotics Platform

The full musculoskeletal model of the virtual rodent controlled by the spinal cord model was simulated in the Neurorobotics Platform (NRP) developed in the Human Brain Project (Falotico et al., 2017). The main components of the NRP are a world simulator, a brain simulator and the mechanism that enables the data flow between the two in a closed-loop. The connection between the body and the brain is specified through a domain specific language (Hinkel et al., 2015, 2017), via Python scripts called Transfer Functions. In these scripts the output of devices that read neuronal output data can be processed and passed as input for the virtual body actuators, and vice versa, the sensory information from the virtual body sensors, in this case muscle length data, can be passed to devices that map sensory data to neural input. The brain simulation, which currently is simulating point-neurons, follows closely the paradigm of NEST (Gewaltig and Diesmann, 2007), interfaced through PyNN (Davison et al., 2009). On the other side of the closed loop the world simulator of choice is Gazebo (Koenig and Howard, 2004), extended to support muscle simulation through OpenSim (Millard et al., 2013a), which provides its’ own muscle simulation engine.

#### 3.2.2 Musculoskeletal embodiment

As described earlier the musculoskeletal system comprises of two elements, the skeletal and the muscle system respectively. Here both systems are elaborated a bit more in the context of the experiment. Developing animal skeletal systems is no trivial task. It involves many complex degrees of freedom and physical properties such as mass, center of mass and inertias. To ease this process, NRP has developed a toolkit for Blender (Open source modeling and animation tool) called RobotDesigner (HBPNeurorobotics, 2019). RobotDesigner allows to automate several steps needed to develop skeletal/robot models to be simulated in the NRP. Using the same, currently NRP hosts state-of-the-art a full skeletal model of the mouse consisting of 110 degrees of freedom. More details about the full model will be soon published following the current article. For the current experiment, the mouse skeletal model is reduced in complexity by constraining all the degrees of freedom except the left forelimb. The forelimb consists of four segments and it is further constrained to only have flexion-extension movements, enough to reproduce the passive extension-active retraction experiment on the M-Platform. The different segments and the joints of the forelimb are shown in Fig. 5.

**Figure 5.**
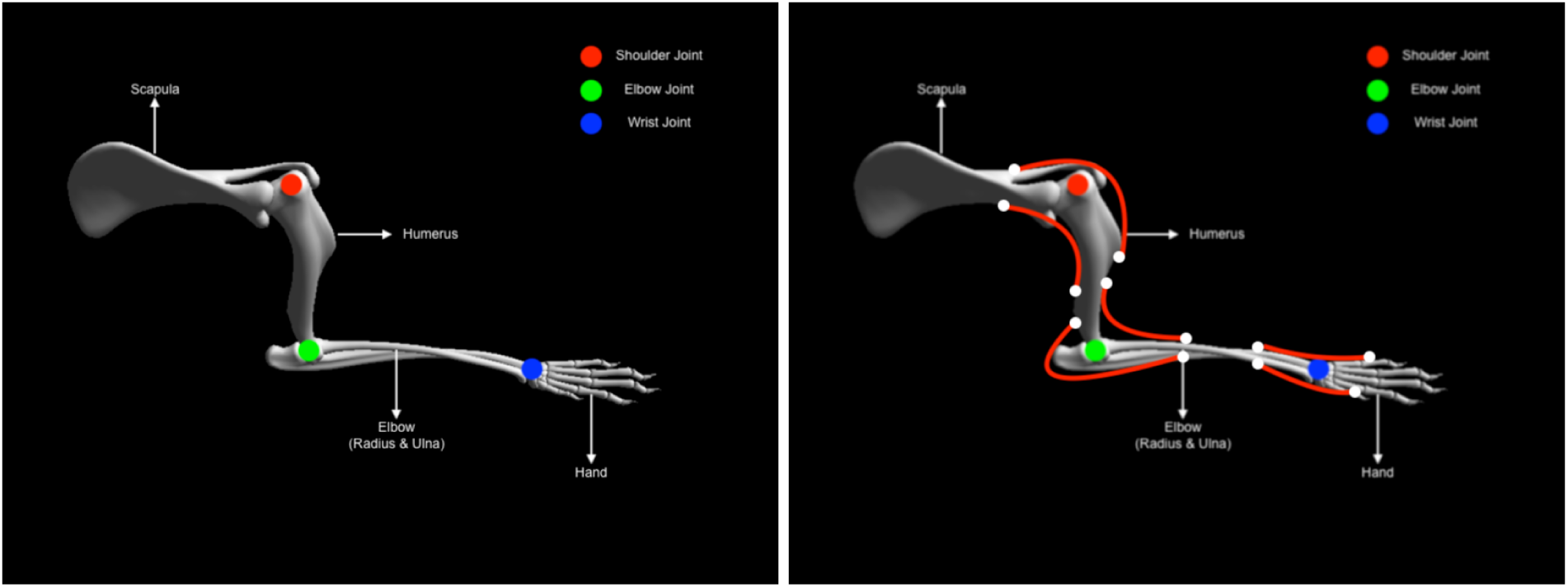
Mouse forearm musculoskeletal system. (left panel) Forelimb skeletal system with three joints (1) shoulder (Ball and socket) (2) elbow (hinge) (3) wrist (hinge). (right panel) Forelimb muscle system with six muscles (1) humerus-extension (2) humerus-flexion (1) elbow-extension (2) elbow-flexion (1) hand-extension (2) hand-flexion

The physical properties of the skeletal system such as mass, center of mass and inertia are automatically estimated based on bounding objects generated for each link (segment) using the RobotDesigner. Once the skeletal system is established, muscles-tendon system can be attached to the bones. As mentioned before, NRP now supports OpenSim for integrating muscle models into physical animal bodies or even robots. In the current experiment a pair of antagonist hill-type muscles were added to each of the joints in the mouse forelimb. The muscle model in OpenSim is taken from Millard and colleagues (Millard et al., 2013b) (see Supplementary Material). Again RobotDesigner offers a unique solution to visualize attachments and easily add muscles to the body in blender. Using the same technique all the muscles for the mouse forelimb were added. Muscle parameters used in the current experiment are hand tuned to produce flexion-extension movements necessary for the experiment. Fig. 5 (right panel) shows the muscle attachments used in the current model.

#### 3.2.3 Robotic rehabilitation platform model

In the real experiment, the mouse forelimb is attached to the sliding mechanism, which is a prismatic joint, driven by a DC motor whose rotational motion is converted into a linear one. The motor that is controlled with a PID controller, whose reference can be set to a position of the joint between the minimum and maximum positions. The controller is enabled when the operator decides to replace the sled in its starting position and is disabled afterwards, so that the mouse can actually pull the sled. In simulation, the same configuration has been implemented. The musculoskeletal mouse forelimb was attached to a simulated M-platform, which has been modelled as a prismatic joint, controlled with a PID controller whose output is directly applied as a simulated force on the joint, assuming ideal actuator transfer behavior. Again, the reference to the PID controller is the position of the prismatic joint in its range, this time normalized between 0 (minimum) and 1 (maximum). To simulate the intervention of the operator that puts the slide back we employed a state machine that automatically controls the slide, by making use of the PID controller setting 1 as a reference. Inputs to this state machine are a list of times at which the slide should be put back. Conversely, to simulate the minimum amount of force that is required to move the slide in the real setup, we deactivated the PID controller in simulation only after a certain activation of the simulated muscles was reached (0.95).

### 3.3 Stroke Models

#### 3.3.1 Brain Network Model with Kuramoto oscillators

To simulate the functional network reorganization during stroke and recovery given by the phase coherence of the macroscopic brain activity reflected in calcium signals, we built our BNM based on Kuramoto oscillators for the local oscillatory dynamics and the AMBA connectome that dictates the strength of the couplings between brain regions (Melozzi et al., 2017; Choi and Mihalas, 2019), Fig. 6, and has been validated with empirical functional data that justifies its use (Melozzi et al., 2019). The AMBA contains 86 cortical regions (43 per hemisphere), of which 18 were included in the field-of-view (Fig. 6, bottom left). The average calcium signal of the pixels entirely located within a brain region was used to represent their mean neuronal activity.

**Figure 6.**
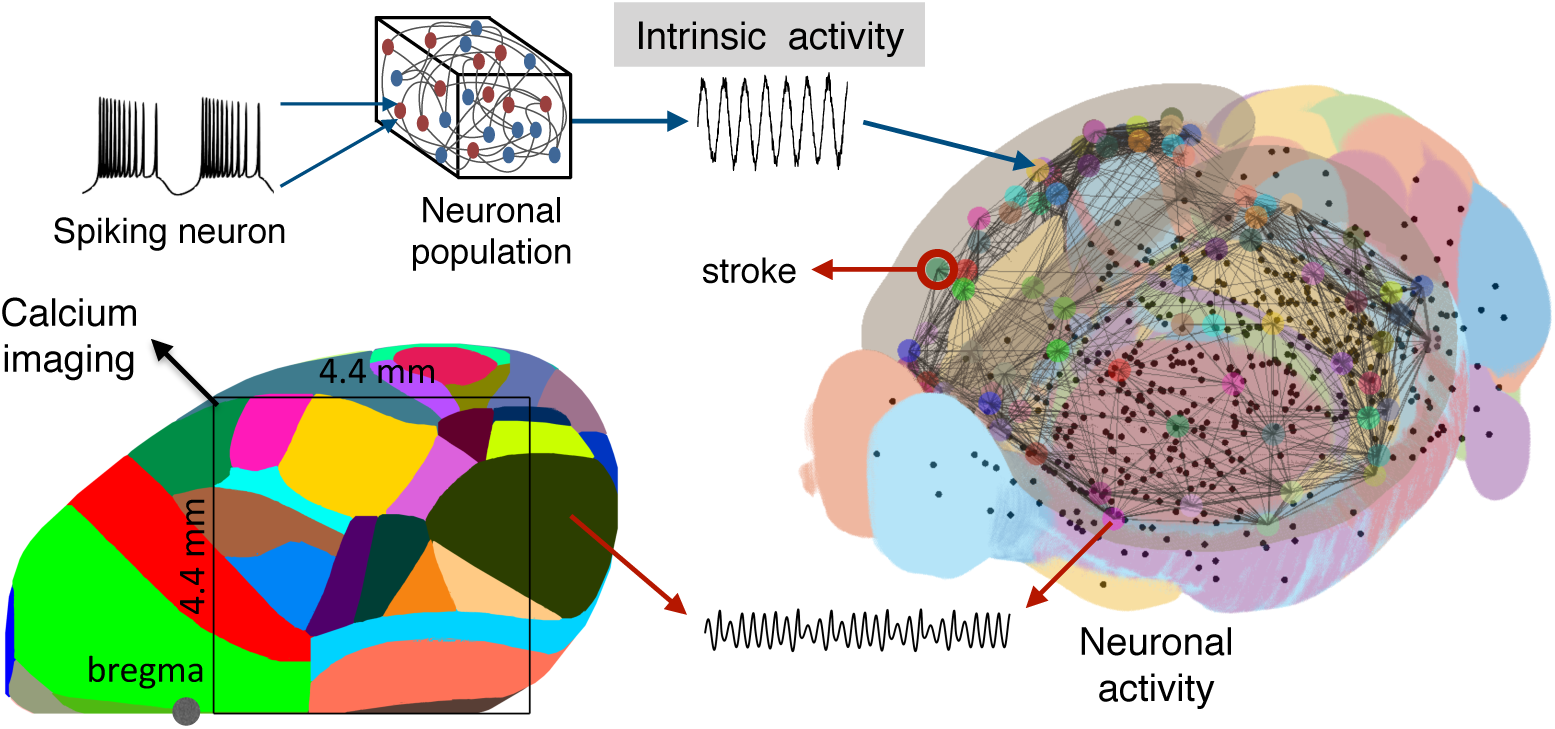
Scheme of the mouse BNM. Brain Network Model consisting of neural masses superimposed over the AMBA connectome simulates the recorded calcium activity. The average oscillatory neuronal activity of the brain regions is described by Kuramoto oscillators, which are coupled due to the fiber tracts, giving rise to the simulated recordings. The brain network (right) is reconstructed from the AMBA, with the centers of subcortical regions being small black dots, while larger the circles are for the cortical regions, with the region of the stroke highlighted. On the left, the field of view during the recordings is overlayed on the reconstructed brain, and different colors represent the cortical regions according to the AMBA.

Besides their simplicity, phase models exhibit rich dynamics and a direct link to more complex biophysical models, while admitting analytic approaches (Roy et al., 2011; Sheppard et al., 2013; Ton et al., 2014; Stankovski et al., 2016). KM (Kuramoto, 1984), as a phenomenological model for emergent group dynamics of weakly coupled oscillators (Pikovsky et al., 2001) is well suited for assessing how the connectome governs the brain oscillatory dynamics that can be reflected in different neuroimaging modalities (Schmidt et al., 2014; Váša et al., 2015; Cabral et al., 2017; Petkoski et al., 2018). The constructed BNM is thus used to identify the structural alterations due to the stroke and the subsequent recovery, using their causal effects on the functional changes captured by the calcium recordings of the cortical brain activity.

Even though delayed interactions due to axonal transmission can be of crucial importance for the observed dynamics of the oscillatory systems (Ghosh et al., 2008; Petkoski et al., 2016, 2018), the impact of these delays is much less pronounced for low frequencies compared with them, as it is the case here. Moreover, the tracing used for obtaining the AMBA Connectome (Oh et al., 2014) does not allow tracking the length of the white fibers. Hence, we assume instantaneous couplings and the utilized model gives the following evolution of the phases for each of the *N* brain regions

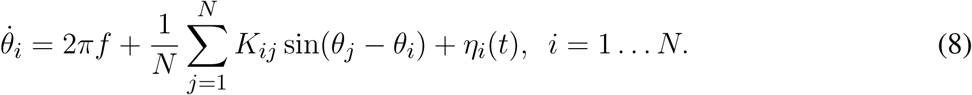

Here the dynamics of each region *i* is driven by the natural frequencies *f* that are assumed to be identical across the brain. A stochastic variability is introduced with additive Gaussian noise defined as ⟨*η*_*i*_(*t*) ⟩ = 0 and ⟨*η*_*i*_(*t*)*η*_*j*_(*t*^*’*^) ⟩ = 2*Dδ*(*t − t*^*’*^)*δ*_*i,j*_, where *D* is the noise strength and ⟨· ⟩denotes time-averaging. The activity of the BNM is then constrained by the structural connectivity, which for every region is represented by the inputs that they receive from the other regions *j* through the coupling strength *K*_*ij*_ = *Kw*_*ij*_. This contains the structural weight of the connectome between these areas, *w*_*ij*_, scaled with the same global coupling *K* for every link.

#### 3.3.2 Spiking network model for simulation of slow wave activity in peri-infarct cortex

Besides the phenomenological neural mass model for the oscillatory activity that we have used in the BNM, we also show an alternative spiking neural model to reproduce local brain activity in the acute phase after stroke. In future, this model should be integrated in the BNM and therefore in the *Embodied brain* closed-loop simulation, either by deriving its mean-field representation, e.g. see (Zerlaut et al., 2017), or by co-simulation. This model aims to reproduce the two-photon calcium signals of a population of spiking neurons located at the peri-infarct area, since it is known that slow frequency patterns of synchronized activity emerge from the damaged areas after an ischemic stroke (Butz et al., 2004; Carmichael and Chesselet, 2002; Rijsdijk et al., 2008; Rabiller et al., 2015).

##### Network of adaptive exponential integrate and fire (adex) neurons

The network consists of an excitatory (regular spiking, RS) and and inhibitory (fast spiking, FS) population of neurons (Fig. 7A). All cells are modeled as adex neurons, which can be described by the following equations:

**Figure 7.**
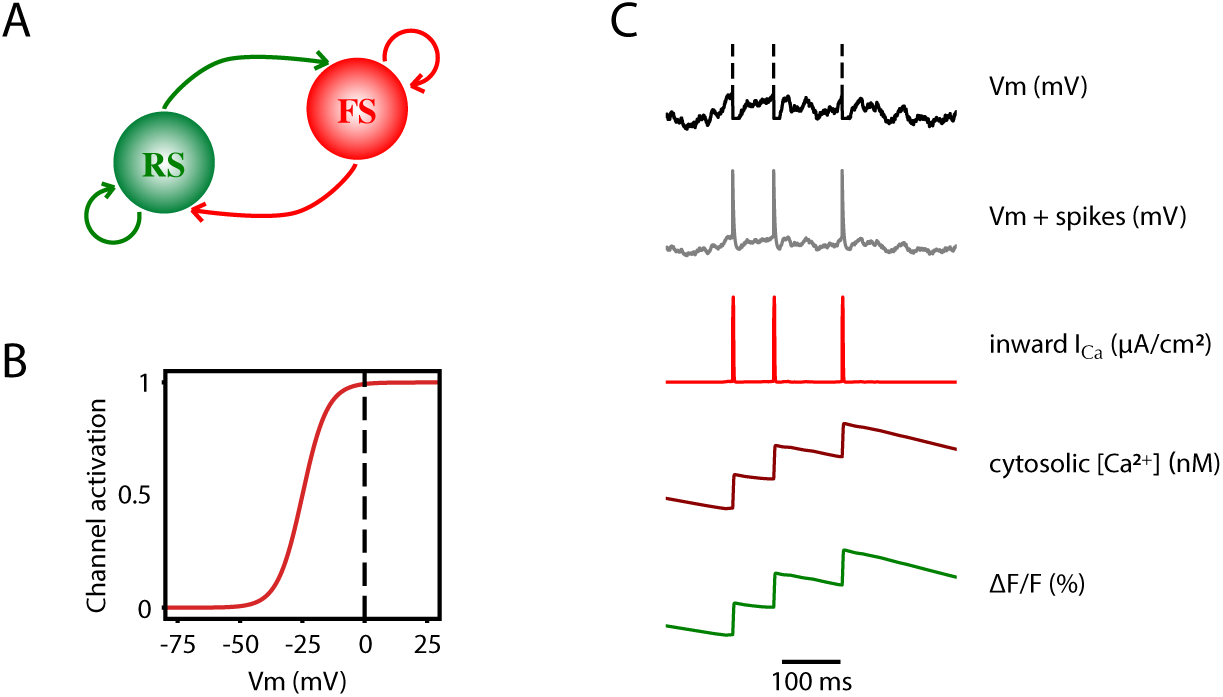
Two-photon calcium signal model from a spiking network model. (A) Schematic connectivity between the excitatory population of regular spiking (RS) neurons and the inhibitory population of fast spiking (FS) neurons of the modelled cortical network. (B) Activation curve of a high-voltage activated calcium channel that is used to compute the inward calcium current (*I*_*Ca*_) from the changes in the *V*_*m*_. (C) From top to bottom, simulated membrane potential of a neuron emitting three spikes which are represented by dashed lines, membrane potential with reconstructed spikes, inward calcium current associated with changes in the membrane potential, cytosolic calcium concentration and fluorescence emitted by the calcium indicator due to the intracellular concentration of calcium.

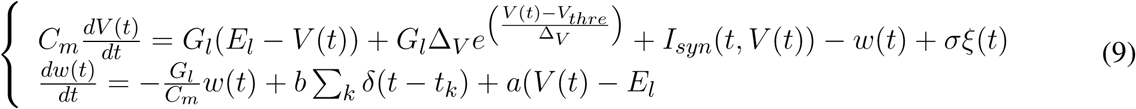

where the synaptic input *I*_*syn*_ is defined as

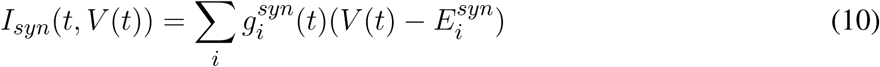

with

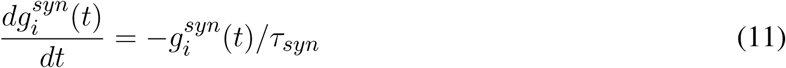

Here, *Gl* = 10 *nS* is the leak conductance and *Cm* = 150 *pF* is the membrane capacitance. The resting potential, *El*, is −60 *mV* or −65 *mV*, for excitatory or inhibitory cells, respectively. Similarly, the steepness of the exponential approach to threshold, Δ_*V*_ is 2.0 *mV* or 0.5 *mV*, for excitatory or inhibitory cells, respectively. When the membrane potential *V* reaches the threshold, *V*_*thre*_ = −50 *mV*, a spike is emitted and *V* is instantaneously reset and clamped to *V*_*reset*_ = −65 *mV* during a refractory period of *T*_*refrac*_ = 5 *ms*. The membrane potential of excitatory neurons is also affected by the adaptation variable, *w*, with time constant *τ*_*w*_ = 500 *ms*, and the dynamics of adaptation is given by parameter *a* = 4 *nS*. At each spike, *w* is incremented by a value *b*, which regulates the strength of adaptation. *b* = 60*pA* was used to model deep anesthesia, and *b* = 20 *pA* for light anesthesia simulations.

#### From spikes to fluorescence of two photon signal

In order to model the two-photon calcium signal the spikes and the values of the membrane potential *V*_*m*_ (Fig. 7C, grey trace) of each neuron were recorded during the simulation, for each level of adaptation. Increases in the *V*_*m*_ lead to an inward calcium current through voltage-dependent channels. We characterized the L-type high voltage activated calcium current *I*_*Ca*_ (Fig. 7C, red trace) as in Rahmati et al. (2016):

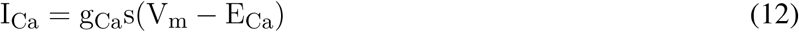

where *E*_*Ca*_ = 120 *mV* and *g*_*Ca*_ = 5 *mS/cm*^2^ are the reversal potential and the maximal conductance of this current, respectively. The steady-state voltage dependent activation of the channel (Fig. 7B), is defined by the Boltzmann function:

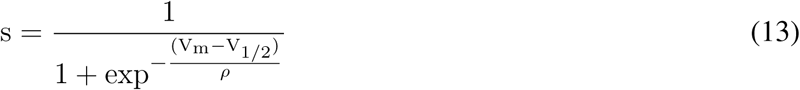

with a half-activation voltage *V*_1/2_ = −25 *mV* and a slope factor *ρ* = 5*mV (Ermentrout, 1998; Helton et al., 2005). The intracellular concentration of calcium (cytosolic [Ca*^2+^], Fig. 7C, dark red trace) increases proportionally to the *I*_*Ca*_ current, and then it slowly decays back to a basal value [*Ca*^2+^]_*i*_ (Traub, 1982):

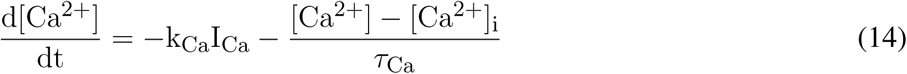

with *K*_*Ca*_ = 0.002 (*nM/ms*)(*µA/cm*^2^)^−1^, *τ*_*Ca*_ = 760 *ms* and [*Ca*^2+^]_*i*_ = 0 *nM*.

Finally, the fluorescence *F* (*t*) associated with the intracellular calcium concentration (Fig. 7C, green trace) is then computed following the equation:

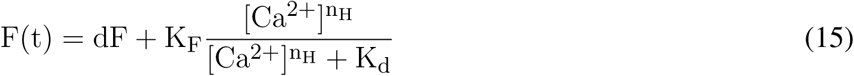

where *dF* = 0 and *K*_*F*_ = 10 are the offset and the scaling of *F* (*t*), *k*_*d*_ = 375 *nM* is the dissociation time constant for GCaMP6f (Chen et al., 2013), a measure of the affinity of the fluorescent indicator to the calcium ion, and *n*_*H*_ = 2.3 is the Hill coefficient (Chen et al., 2013).

## 4 RESULTS

Here we show first the results we obtained on the simulation of goal-directed movements (“Movement-driven models” pipeline) and then on the modeling of brain alterations after stroke (“Stroke models” pipeline).

### 4.1 Simulation of the experiment on goal-directed movements

As the first component of the proposed framework (“Movement-driven models”), we simulated the experiment on goal-directed forelimb pulling in the virtual environment and validated the simulation on experimental data.

In the in-vivo experiment, two healthy mice were trained on the M-Platform to perform active pulling of the forelimb. As we expected, the contralateral motor cortex showed a highly coherent activation with the kinetic data. The coherence between the force applied by the animal and the signal recorded in the CFA was evident both in the low and in the high frequencies band (Fig. 8). For the data that were later used in simulations we focused on the high band (300 to 40k Hz); in particular we found an high activation of the motor cortex around the force peaks for both multi-unit activity and single units analysis. This result proves that for each recording the SUs were successfully extracted by the multi-units. The PSTHs was used to evaluate the temporal behaviour-related spike activity of every single unit. The behaviour around the force peaks was different according to the single units selected, but all of them showed that the activity began to increase before onset of force peaks and came back to the resting value after 0.4 s from the onset. In order to simulate the descending signal from the motor cortex generating the movement of the forelimb, we employed these neurophysiological recordings. In particular, the events resulting from the single unit spike sorting were given as spike times for static spike generator in the neural simulation. As the number of recorded neurons was low, the spike generators were copied 100 times, while also adding gaussian noise (with mean = 0ms and standard deviation = 5ms) to the spike times of the copies to avoid synchronicity. As the neural recording originates mainly from neurons that control the pulling, we decided to connect the descending stimuli only to interneural populations associated with muscles that are active during the pulling, i.e the flexors of the the two actuated joints. Therefore, the antagonistic muscles would only actuate thanks to spinal reflexes. To tune the parameters of these connections, and to produce a muscular activation that was similar in amplitude to the force recorded in the in-vivo experiments, we performed a preliminary set of experiments, without the simulated embodiment, in which we empirically tuned the synaptic weights and number of connections. Due to the absence of the embodiment, at this stage there is no muscle spindle activity and thus no sensory feedback enabling reflexes. Then, the spinal cord model described in Section 3.1 was connected to the mouse forelimb. In principle, the musculoskeletal embodiment has three pairs of muscles, but the one controlling the paw is not significantly involved in the pulling of the limb. We did not consider those when building the neural network to decrease simulation times. Thus, we replicated the same spinal cord circuitry two times and connected it to the four muscles controlling the elbow and shoulder joints, named <u>humerus</u> and <u>radius</u> in the simulation. In the closed loop simulated by the Neurorobotics Platform, the output of the spinal cord model (muscles activation between 0 and 1) could be directly given to the simulated mouse actuators, while the muscle lengths and contraction speed had to be normalized before sending them to the muscle spindles models.

**Figure 8.**
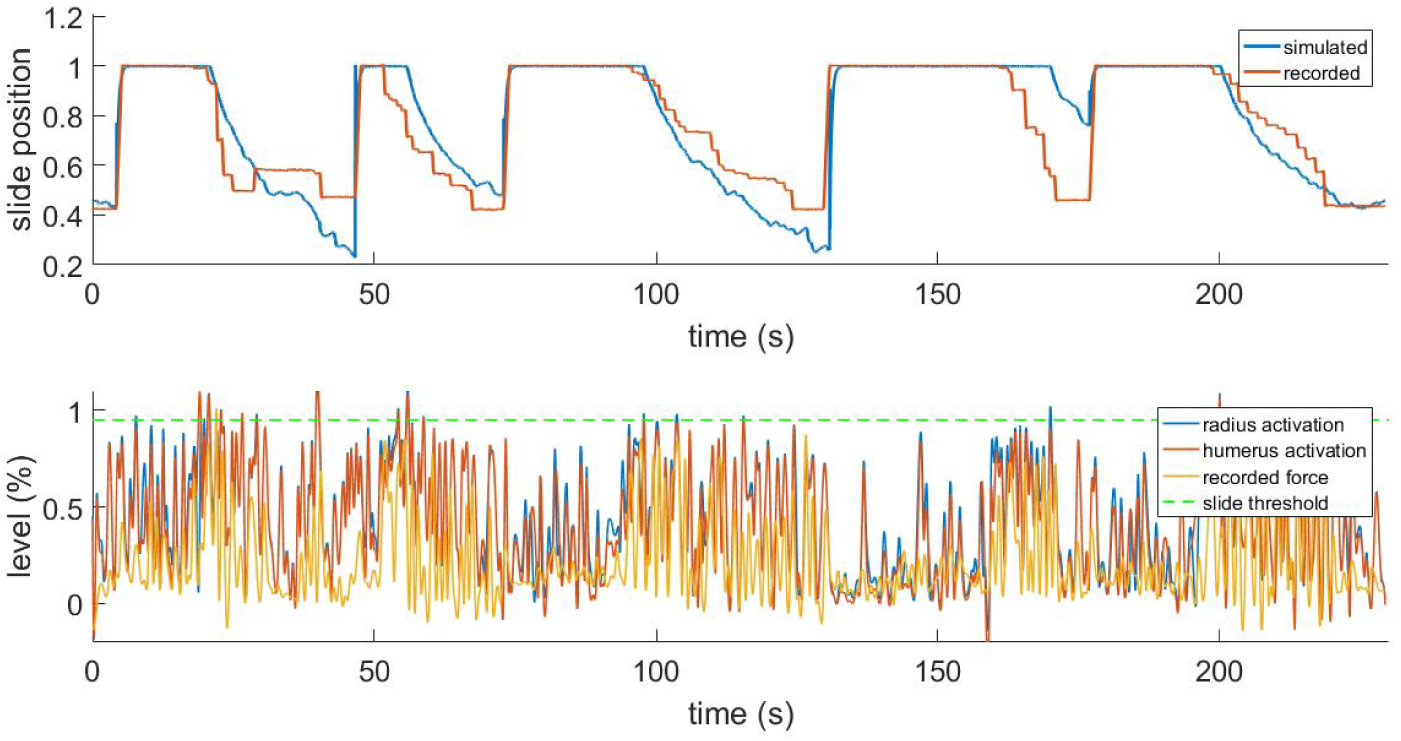
Results from the simulated pulling experiment. Comparison between the simulated slide position and the one recorded in the in-vivo experiment (top) and comparison between simulated muscle activation levels and force applied to the physical slide (bottom). To increase the readability of the bottom figure, an upper peak envelope was applied to the signals.

In Fig. 8 we show the results for a simulation trial and a comparison with data recorded from a physical experiment. We employed kinematic data recorded alongside neural activity in the same in-vivo experiment: position of the slide and force applied to the slide through the trial. As expected, by comparing the activation levels with the normalized force applied by the mouse to the slide we can observe that the flexor muscles are active when there is also a force recorded, and conversely, there is low activation when the slide is still. It is also worth mentioning that, although the two muscles receive the same inputs from the descending stimuli, their activation levels are different due to the feedback circuitry of the spinal cord and the activity of muscle spindles, which are different for the two muscles. Thus, the output of the spinal cord circuitry is not a mere filtering of the input signals, but it also takes into account the feedback from the embodiment, which can change during the experiment. This effect underlies the importance of embedding neural circuits in a proper, realistic embodiment. The comparison between the simulated slide position and the recorded one shows that, thanks to the recorded neural activity, the muscles are able to overcome the force threshold and release the slide, and that an actual pulling is performed. Every pulling episode in the trial is reproduced, even if with different degrees of accuracy. Overall, the mean absolute error between simulated and recorded slide positions is 13%. The main discrepancies between the simulated and the recorded data come from the fact the in the simulated mode, the muscular activity is mostly directly proportional to the neural activity, while in the recorded data this is not always the case. While there is clearly a correlation between presence of neural activity and motion, the intensity of such activity sometimes does not match the intensity of the motion.

### 4.2 Local and global brain simulations after stroke and rehabilitation

#### 4.2.1 BNM for brain connectivity changes after stroke and rehabilitation

Within the second pipeline of the framework (“Stroke models”), in this subsection we simulated different extents of the brain injury and rehabilitation-induced plasticity after stroke. The results from the simulations are compared with the experimental data, Fig. 2, allowing us to find the best fit with the empirical functional reorganization in the parameters space of the structural changes in the white matter connectivity.

The averaged calcium activity has a local peak in the power spectrum at around 3.5 Hz (Fig. 9, (A)), which is within the band relevant for the resting state activity. We hence focused the analysis of the experimental data on the the upper delta band, 2.5 − 5*Hz*, where we consequently band-pass filter the signals. From these we calculated the pair-wise PLV in each condition, thus constructing the FC matrices for the cortical regions of interest. Finally, to remove one condition, we calculate the changes of the FC during stroke and recovery compared with the healthy state, and this is the data feature that is then compared with the simulated data. For this we use the model described in 3.3.1, to identified which scenarios of structural alterations cause the best agreement with the data in the modelled FC alterations, Fig. 9. To minimize the effect of tissue displacement after stroke (Brown et al., 2007), the analysis includes only 12 ipsilesional regions located outside the stroke core, Fig. 9 (B and D).

**Figure 9.**
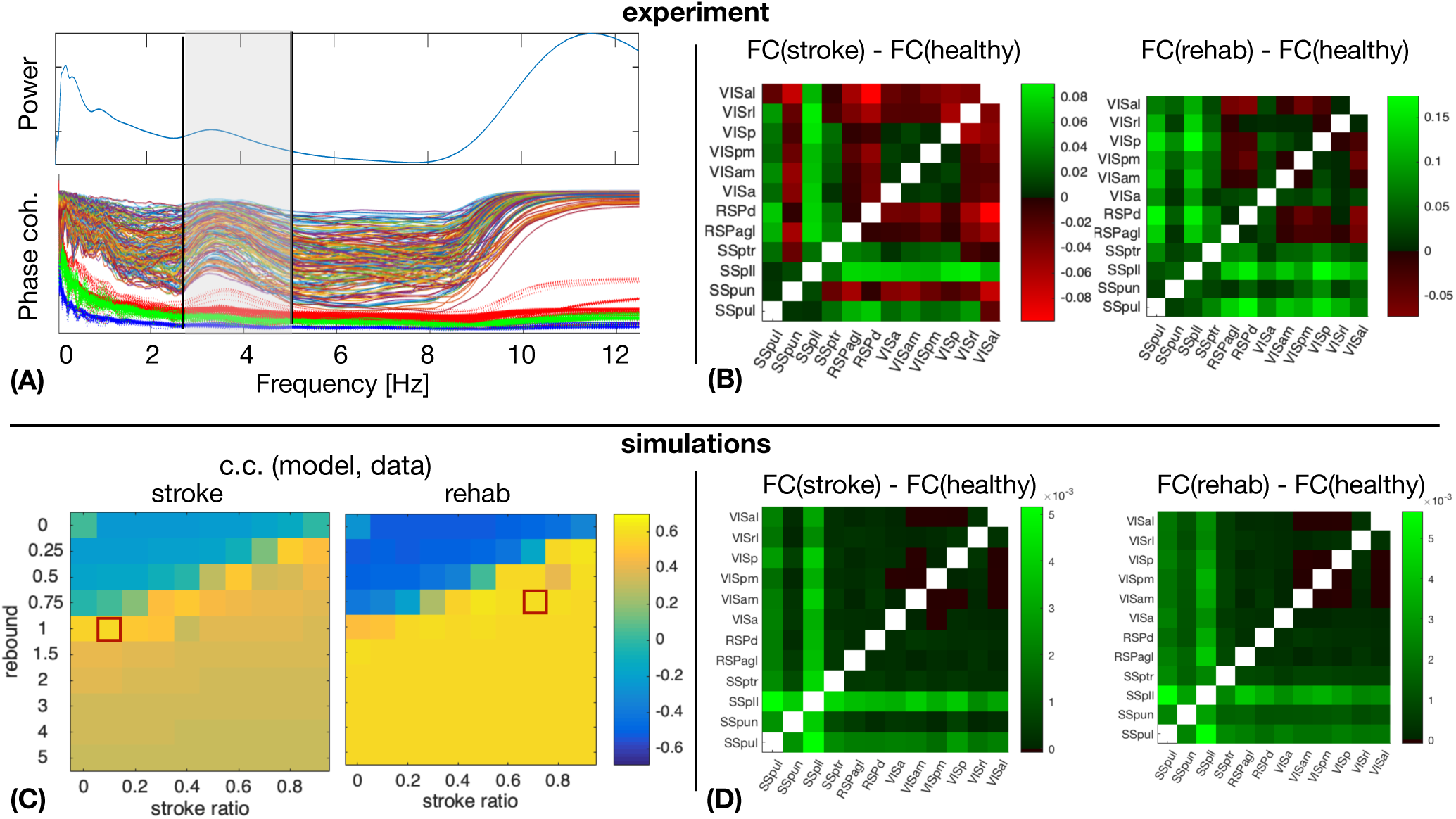
Simulated and empirical Functional connectivity and fitting the model. (A) Average power spectrum across the regions and the PLV values for each pair of regions (thin lines) during healthy state, as well as the significance levels from the surrogates (thick dotted lines). Vertical black lines show the boundaries of the lower and upper *δ* bands. (B) Relative changes of the FC at stroke and at rehabilitation compared to the healthy control for frequency band *f* = 2.5 − 5*Hz*. (C) Cross correlation of the model upper triangles of FC between the model and the data for fixed global coupling *K* = 4.3 and different levels of stroke (0 for complete damage and 0.9 for damage of 10% of the links) and rebound connectivity (0 for no rewiring and 5 for overall rewiring with strength of 5 times of the damaged links). Parameters: frequency *f* = 2*Hz*, noise strength *D* = 1. (D) Simulated relative changes of the FC at stroke and rehabilitation relative to the healthy control for the working points marked with red squares in the parameters space in the panel (C). The abbreviations for the areas in (B) and (D) are: VIS - visual, RSP - retrosplenial, SS - somatosensory, al - anterolateral, rl - rostrolateral, p - primary, pm - posteromedial, am - anteromedial, a - anterior, d - dorsal, agl - lateral agranular part, ptr - primary trunk, pll - primary lower limb, pun - primary unassigned, pul - primary upper limb.

The stroke affects not only the inherent activity of the rM1, but all the connected regions. However, the precise breadth and magnitude of the structural damage, namely which links and to what extent are they disabled over time, is unknown. Similarly, it is not known which new links are created or reinforced during the spontaneous recovery or what is modulated by the rehabilitation. On the other hand, the stroke was shown to consistently change the alignment of dendrites and axons towards the core *in vivo* (Brown et al., 2007), possibly meaning altered SC, confirming previous works on structural rewiring after stroke (Dancause, 2005; Nudo, 2013). Hence a numerical exploration of the different possibilities of the stroke and rewiring in the large-scale BNM is used to unveil the most probable structural alterations associated with stroke and recovery. The calcium activity in the upper delta band that was chosen for the analysis shows highly coherent co-activation of different parts of the cortex, compared with the surrogate time-series, Fig. 9 (A). We compared the functional reorganization associated with spontaneous recovery after stroke (“stroke” group) to rehabilitation-supported recovery (“rehab” group). The changes in the functional connectivity in “stroke” compared to “healthy” mice (Fig. 9 (B), left matrix) indicate an increased co-activation of all but one somatosensory areas in the chronic phase after stroke, while visual areas have increased connectivity with all the regions, and reduced with the retrosplenial cortex. In the rehabilitated mice (Fig. 9 (B), right matrix), the increase in connectivity of the somatosensory is even higher across all the areas, and there is also an increased FC of the visual areas between each other and with the somatosensory regions.

A phenomenological neural mass is used to simulate how ipsilesional FC is changed by stroke and rehabilitation based on the modifications of the SC. For this, we systematically modified the SC to account for various impacts of stroke and subsequent recovery, in order to find the best match with the patterns observed in the data. The damage due to the stroke is assumed to be homogeneous across the links connecting rM1, but their magnitude is varied from 10% to 100%. Similarly, after the recovery it is assumed that 0 to 500% of the lost connectivity due to stroke is restored homogeneously across the regions with preexisting links towards rM1, proportionally to the initial strength of their link to rM1. We thus explore the possibility of up to 5 times of weights of the damaged links to be redistributed along the rest of the links of the nodes directly connected with the infarct area, in order to also allow for over-compensation of the lost direct connectivity. The absence of time-delays and the focus on the phase locking, makes the model insensitive on the chosen frequencies (Petkoski et al., 2016), which are therefore fixed in the simulations. The natural time-variability of parameters Petkoski and Stefanovska (2012) is assumed to be stochastic (Petkoski et al., 2018). We hence fix the level of the noise and we explore the impact of the global coupling *K* and the described strategies of the stroke and recovery. For each combination of parameters we obtain the same metric of FC as for the empirical data. The parameters space for the agreement between the modelled and the experimental data about the changes in the FC for the two parameters of the stroke-induced structural changes are shown in Fig. 9 (C).

Panel (D) of Fig. 9 illustrates the simulated FC for spontaneously recovered “stroke” and “rehab” mice compared with pre-stroke conditions (“healthy” group), for fixed global coupling and for points in the parameters space of the stroke damage and rebound connectivity that show the best fitting with the empirical data. Comparing the simulated, Fig. 9 (B), with the empirical FC, Fig. 9 (D), we see that the best agreement is achieved for the FC of the somatosensory areas, while that of the visual cortex areas could be improved by testing different damage and rewiring strategies for those regions. From the model fitting for different parameters, it is also visible that generally better fit is achieved if the extent of damaged links is decreased due to rehabilitation-induced remapping. There is also a similar tendency for the rebound connectivity to be decreased due to recovery training, although there are other possible recovery paths that keep roughly the same level of rebound connectivity. In conclusion, the systematic exploration of the model parameters to best fit the empirical data, allows us to obtain the sufficient structural changes that can reproduce the modulation in FC after stroke and rehabilitation.

#### 4.2.2 Simulation of the calcium activity of the peri-infarct network after stroke

Stroke profoundly alters the functionality at the local level in addition to long-range connections. The local network next to the stroke core switches to slow wave activity (Butz et al., 2004), a type of brain oscillation that is observed during deep sleep, but also during anesthesia and other pathological brain states (Sanchez-Vives et al., 2017). Understanding the changes in the activity patterns at the level of the peri-stroke region is necessary to get insight on the possible mechanisms that underlie functional recovery. In order to explore the mechanisms that drive the neuronal networks of the peri-stroke areas to oscillate, we developed a model that reproduces the spiking activity of a local network during slow oscillations and extended the model to provide the two-photon calcium signal that one would record from that network. We compared the simulated calcium data with that of the two-photon experiments conducted in anesthetized mice (see the “Stroke models” box in Fig. 2).

We propose that a deficit in neuromodulation produced by the decreased cerebral blood flow in the periphery of the region affected by the stroke could be responsible for the emergence of slow oscillations and the general flattening of the EEG through an increase on the level of adaptation of the neurons (Nghiem et al., 2020). To test this hypothesis, we developed a spiking network model capable of reproducing the spontaneous activity of a cortical network during different depths of anesthesia (Fig. 7). The strength of adaptation in the model can be regulated to produce different types of oscillations. In particular, we aimed at reproducing the changes in frequency of the slow oscillation observed both in the two-photon and in the wide field calcium imaging *in vivo* experiments when the anesthesia is reduced (Fig.10A). When adaptation is strong, the model of deep anesthesia produces slow oscillations at 2.1 Hz, while decreasing the strength of adaptation (model of light anesthesia) leads to slightly faster slow oscillations, at a frequency of 2.31 Hz (Fig. 10B). Thus, we show that varying the strength of adaptation allows to reproduce the increase in frequency of the slow oscillations observed in the calcium imaging experiments when decreasing the level of anesthesia.

**Figure 10.**
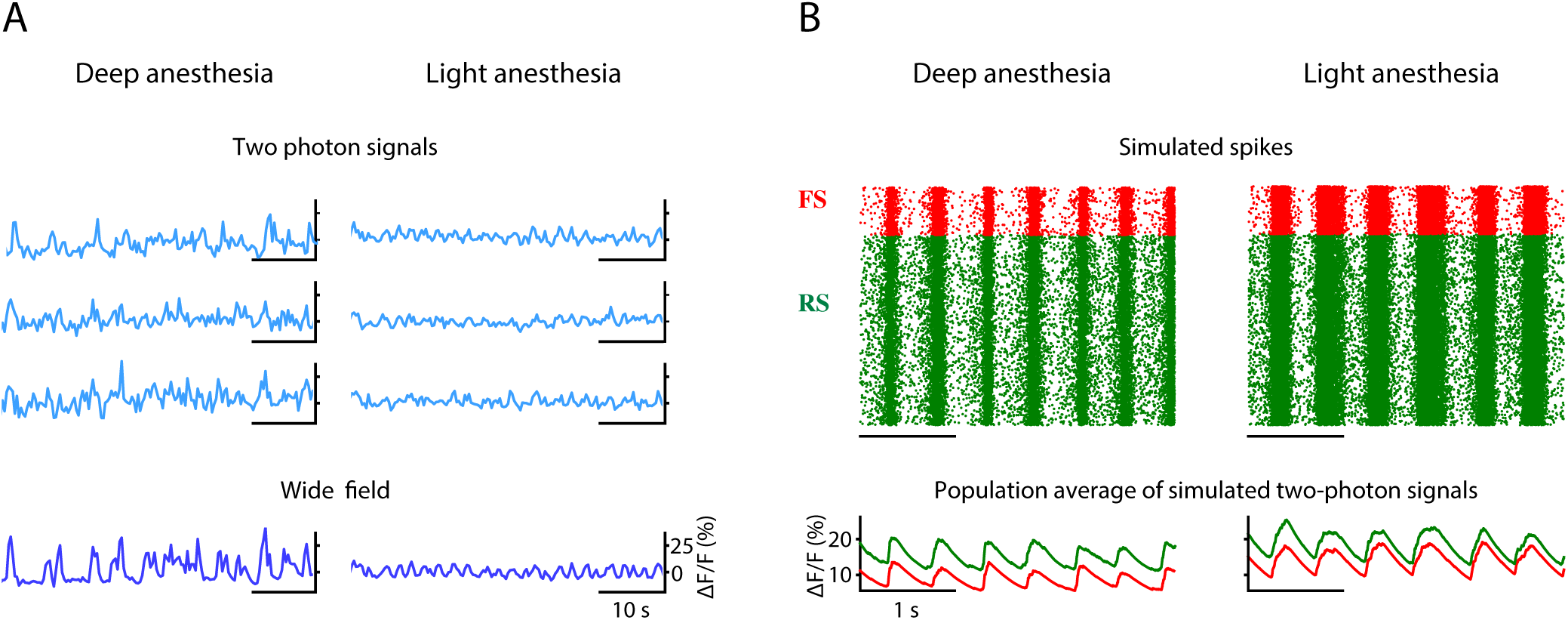
Simulation of peri-stroke local network oscillations: experiments and models. (A) Fluorescence traces obtained from three example cells (two-photon signals) and from the entire field-of-view (wide field signals) recorded in mice under deep and light anesthesia. (B) Raster plot of the spikes (top) and averaged two-photon calcium signals (bottom) computed for the inhibitory (FS, in red) and excitatory (RS, in green) populations in a model of deep or light anesthesia.

## 5 DISCUSSION

In this paper, we explored the steps and methods that are needed to develop a simulation model of a complete experiment. We designed, validated and combined many components, even though they are obviously not exhaustive to encounter for the complexity of the real world.

We first simulated execution of goal-directed movements with displacement of objects in the virtual reality setting. Results show that we could replicate with high accuracy the displacement of an object by a virtual (healthy) mouse in the simulated environment (Fig. 2). While this simulation is only an approximation of brain-body interactions, it showcases the capability to simulate large scale neural networks as well as body dynamics in a virtual environment.

The second pipeline of the framework involved modeling brain injury. We validated the potential of a brain network model to predict the long-range stroke-induced connectivity changes measured in a real experiment. We also tested an oscillatory spiking network model to simulate local peri-infarct activity after stroke. In addition, this model could simulate the fluctuation in calcium concentration due to spiking activity of homogeneous neuronal network, thus allowing modeling of calcium imaging data.

Towards the mechanistic understanding of behavior, few studies already provide tools for closed loop neuroscience (Weidel et al. (2016); Mulas et al. (2010); Tessadori et al. (2012), for a review see Potter et al. (2014)). In addition, recent studies took advantage of virtual reality (VR) experiments conducted under controlled environment, where behavioral strategies could be isolated and tested (Dombeck and Reiser, 2012). In a VR experiment, a simulated environment is updated based on the animal’s actions (Chronis et al., 2007; Ritter et al., 2001; Reiser and Dickinson, 2008; Dombeck et al., 2010). The main drawback of this approach is that the activity of animals dictates not only the response of the VR but also the properties of the neurons being measured. As a consequence, the closed-loop VR system shall then be optimized on-line based on the animal’s behaviour, which is very challenging. The approach we propose here instead is based on an off-line simulation, that allows exploring multiple dimensions in the parameter space of the dynamical model of mouse brain and the environment. Anyway, both strategies are synergistic with the research of effective functional brain machine interfaces (Santhanam et al. (2006)).

### 5.1 Movement-driven models Closed-loop

The results showed in Section 4.1 demonstrate that is possible to achieve realistic simulations by integrating some of the components described previously. Accuracy of the closed-loop simulation could be increased by removing some simplifications that are currently in place. Some of them are related to the physical models of the slide and of the musculoskeletal embodiment. Regarding the former, a more accurate slide simulation will allow to introduce friction effects that are occurring in the real setup, thus we could avoid putting a muscle activation level threshold for the release of the slide. Moreover, more detailed spinal cord and musculoskeletal models will be essential to simulate finer movements.

Results shown in Fig. 8 demonstrate the the system is able to simulate the pulling task, albeit with some inaccuracies on some pulling trials. A presumable cause of this inaccuracy can be identified in the low number of neurons (less than 20) that is possible to record during an experiment on the platform with the 16 channels linear probe. For this reason, it is possible that the selected units do not encompass the entire population of neurons involved in the movement. This issue could be mitigated by employing a multi-unit analysis, however, this will add to the inputs a significant background activity which may not be useful to generate the pulling movement.

Many parameters of the spinal cord circuitry can be adjusted, depending on the inputs, to accurately reproduce the movements recorded in the in-vivo experiments. While in this work the tuning was done manually, a more effective and generalized way would be to use different recordings, both neurophysiological and kinematic, and employ an optimization similar to what has been done in (Sreenivasa et al., 2016).

The level of detail of the spinal cord circuitry can clearly be improved. In this work we modelled a minimal set of components that were capable of replicating experimental data with a certain degree of realism. To achieve this, it was decided not to arbitrarily increase the complexity of the models by adding subcircuits whose impact cannot be clearly measured from a comparison with experimental data. Among these, it is worth mentioning the inclusion of proprioceptive feedback from Golgi tendon organs, which could be potentially implemented with computational models such as (Mileusnic and Loeb, 2006) or the one already included in a spinal cord model in (Mugge et al. (2010)). Perhaps more interesting is the modulation of muscle spindle sensitivity from *γ*-motoneurons, as this is crucial in the control of both voluntary and involuntary movements. While including a population of *γ*-motoneurons could be done by replicating populations of *α*-motoneurons, measuring the impact of adding this component is not trivial, especially considered that there is no experimental data measured, in the rehabilitation setup, that can be used to validate the addition. As such, we decided not include *γ*-motoneurons in the spinal cord circuitry.

### 5.2 Stroke models Closed-loop

AMBA was previously tested and demonstrated to have a predictive value for the resting state dynamics in healthy conditions, compared with the gold standard individualized diffusion tensor imaging connectome (Melozzi et al., 2019). One of the main aims of the stroke modeling pipeline in this study is to validate the use of AMBA in the cases when there are significant changes in SC as compared to the healthy state for which it was obtained (Oh et al., 2014). This requires finding the most probable structural alterations corresponding to the stroke and recovery. From the perspective of the integrative neuroscience, this is especially important as it will allow further application of these altered connectomes validated from the resting state FC, to generate the particular brain dynamics associated with active forelimb pulling on the M-platform by stroke and rehabilitated mice.

To this aim, the present study suggests that rehabilitative training could reinforce the connectivity between motor and visual areas. The iterative loop between experiments and modeling goes towards the confirmation of this hypothesis via stimulation experiments. New experiments shall verify the necessity of this feature in promoting the recovery by stimulating the connections between motor and visual cortex, and modulation of FC could be achieved via optogenetic stimulation, which recently showed to be a promising approach in stroke recovery (e.g. Pendharkar et al. (2016); Conti et al. (2020); Cheng et al. (2014)).

The results from the model identify routes from the stroke to the recovery in the parameter space that can be related to neurophysiological quantities, such as the white matter tracts. We could thus determine links that need to be restored, or prevented from being established, for a successful recovery. One such a recovery path proposed by the model is the rebound in SC after rehabilitative training, and this is especially true for the links involving the visual-associated areas. The proposed rebound is due to newly established links from the regions afferent to the site of the stroke. This can lead to overall overcompensation for the SC, and some of these scenarios could be possible paths for recovery. However, it remains to be seen whether the structural changes of such magnitude can be achieved. Several studies previously showed that axonal growth is stimulated by neurorehabilitative activities after stroke, and that sprouting can extend to widespread brain systems (Carmichael et al., 2017). New experiments aimed at verifying the SC modifications shall verify the hypothesis on the importance of modified connectivity in the visual areas for recovery. In addition, stimulation experiments can also strengthen certain links, and with our modelling framework we can virtually compare the effects of each such modification to the observed dynamics.

For the best fitting of the data one would also need to allow different levels of the global coupling, which governs the global level of synchronization and that is already shown to be increased during stroke (Falcon et al., 2016; Corbetta et al., 2018), thus decreasing integration and information capacity (Adhikari et al., 2017) and modularity (Falcon et al., 2015). Thus, one could more precisely identify the path from stroke to recovery for a wider parameters range. This also includes numerically testing different scenarios for heterogeneous connectivity reinforcing (Nudo, 2013) such as reinforcing of contralateral links in general, or those to contralateral stroke region only, or towards the nodes (ipsi-, contra-lateral, or both) that were connected to the damaged region prior stroke.

Possible problems could arise from the alignment of the experimental data, especially after the stroke, due to the shrinkage and the movement of the tissue (Brown et al., 2007; Allegra Mascaro et al., 2019). We have tried to avoid this by excluding from the analysis the regions adjacent to the stroke, but this reduces the predictive value of the model due to smaller number of analyzed regions.

Finally, these experiments provide a picture of the ipsilesional functionality after stroke and rehabilitation, but many other regions are involved, including the contralesional hemisphere (see, for instance, Dodd et al. (2017)). In the next experiments, the focus shall be on recording with a higher sampling rate to capture wider spectrum of brain dynamics, and on enlarging the field-of-view of the wide-field imaging setup to provide longitudinal pictures of cortical functionality over both hemispheres. The latter should also refine the fitting across parameters, which now contains large areas or similar level of predictability, thus offering more precise recovery path. Individualized connectome data by Diffusion Tensor Imaging during the recovery process is another aspect of the future experiments that should test the predicted changes in the structure that we propose to be the cause of the observed functional alterations of different conditions. In addition, higher resolution SC performed with light-sheet microscopy on individual mice (Allegra Mascaro et al., 2015) could test the model prediction at the final time point of the experiment. As a final step, an individualized therapy could be proposed targeting specific parts of the brain (Spalletti et al. (2017), Allegra Mascaro et al. (2019) and Conti et al. (2020)), depending on the location and the size of the stroke.

### 5.3 Integration

We propose viable strategies to integrate the brain models described here and to embed them within the *Embodied brain* framework on the NRP (Falotico et al. (2017)) (pictured by the overlapping green and red boxes in Fig. 11). Before applying it to the simulation of the whole-brain dynamics, the spiking neurons model shall be extended to include the heterogeneous long-range connections either via mean-field approximation or by means of co-simulation with other neural masses (see the spiking neurons model that receives calcium imaging data in Fig. 11). In addition, a model for embedding spiking model modules into the whole brain model is currently under development (displayed as a gray arrow from the spiking neurons to BNM in Fig. 11). This work includes validating neuronal mass models against high-dimensional neuronal networks. Once available, this tool will allow bridging the scales of brain models with different levels of description, and they will be then implemented in the NRP and integrated into the *Embodied brain* framework (grey arrows in upper box of Fig. 11).

**Figure 11.**
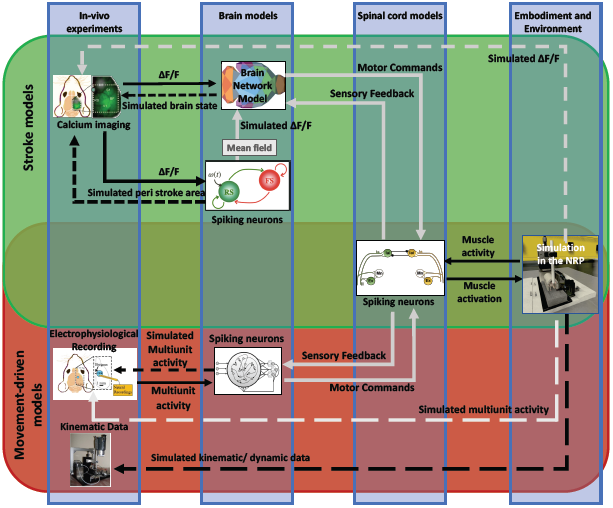
Future perspective of data and simulations. The scheme depicts the approach to build the framework from data to models and back. The workflow from data to models and simulation in the *Embodied brain* closed loops is shown. The upper, green box shows the *Stroke models* closed loop, the lower, red box shows the *Movement-driven models* closed loop. Colored images represent experiment data, brain and spinal cord models, and simulation of the environment (from left to right). Connections between the components are presented as arrows: solid lines represent the output provided to other blocks; dashed lines indicate the output data of the models that are used for comparison with real data for validation. In gray, models and connections that are still under development. The overlapping green and red region pictures the future integration of the two pipelines, and in particular of the brain models within the NRP.

To integrate the large-scale BNM with the proposed spinal cord model, we propose to modulate the activity of the spiking neurons in the spinal cord by the output of the cortical regions, mainly those related to the motor activity (displayed by grey arrows in the *Stroke models* closed loop, green upper box in Fig. 11). In particular, the firing rate of the neurons in the spinal cord that triggers the movements on the NRP can be driven by the mean activity of the cortical motor regions, or by some specific patterns of their co-activation, such as a high-level activity propagation, similar as the one observed during the movements. In this way, the mean neuronal activity of the brain regions at different conditions would trigger movements at the NRP using the activity of the spinal cord. For the feedback link of the sensory activity, we envision the information about muscle activity and limb displacement, which is encoded into the firing patterns of the spinal cord spiking neurons, to directly modulate the mean activity of the sensory motor regions (displayed as a dashed grey arrow in the *Stroke models* closed loop, upper box in Fig. 11). This on the other hand would impact the overall brain network dynamics, including the activation patterns of the motor regions.

To allow the flow of information from the brain to the virtual environment, we anticipate that the next step will be the integration of a spiking network model of motor areas upstream to the spinal cord model. This data-driven model of the motor cortex will include populations of pyramidal neurons and interneurons that can be functionally attached to different lower circuits (displayed by grey arrows in the *Movement-driven* closed loop, red lower box in Fig. 11). This integration in the proposed framework can be an effective strategy to effectively close the *Embodied brain* loop.

## 6 CONCLUSIONS

To summarize, in this study we proposed a methodological framework (named *Embodied brain*) to investigate a “brain in the loop” by a constructive refinement of experiments and simulation of an embodied mouse.

Our findings suggest that simulation of real experiments within the proposed framework will help better understand the complex mechanism that underlies the generation of behavior. Nevertheless, the actual advantages of the “Embodiement” approach, still under construction, are largely unexplored. Even though some aspects of complex animal behavior may be represented with good accuracy by modelling single neural components, without embedding the neural simulations in a physical embodiment it is impossible to show the effect of such neural systems on the body and the surrounding environment. In our study, it would be impossible to assess whether or not the neural models are capable of performing the pulling task with any degree of accuracy, computed on the kinematic data. Furthermore, we believe that new features (e.g. activation of different brain regions for performing the same task due to degeneracy (Price and Friston, 2002) and its impact for stroke and recovery) will be disclosed by the simulation of the entire experiment. In conclusion, the framework shown in this study will advance the field by formulating new hypothesis on the mechanism underlying goal-directed voluntary movements, to be validated on *ad hoc* designed experiments.In general, the framework could simulate new types of experiments that cannot be run in the real word. Last but not least, the virtual environment will be an essential tool to reduce the number of animals used in the experiments, thus making the “Reduction” rule on animal experimentation a feasible goal.

## AUTHOR CONTRIBUTIONS

ALAM, EF, SP, LV, MOG, FSP conceived the study; ALAM, MP, NTC, EC, FR, CS performed experiments; ALAM, EG, SP, LV, MP, NTC, EC, STR, EA, CHB, TBL wrote the paper; All authors agreed with the manuscript; SP, VJ, NTC and AD developed the brain models; LV and EF developed the spinal cord model; LV, AVA, EA, ST developed the simulation in the NRP.

## ACKNOWLEDGMENTS

We thank Krister Andersson, Oliver Schmidt, Martin Øvsthus, Ingrid Reiten and Jan G. Bjaalie of the Human Brain Project curation team for expert assistance to share data via the Human Brain Project Neuroinformatics Platform.

## DATA AVAILABILITY STATEMENT

The datasets and data description on wide-field calcium imaging generated for this study can be found here: Allegra Mascaro, A., Conti, E., Sacconi, L., & Pavone, F. (2019). Fluorescence cortical recording of mouse activity after stroke [Data set]. Human Brain Project Neuroinformatics Platform. DOI: 10.25493/Z9J0- ZZQ.

The raw datasets on neurophysiological recordings for this study can be found here: Pasquini, Maria, Spalletti, Cristina, Caleo, Matteo, and Micera, Silvestro. (2019). Recordings of Caudal Forelimb Area in healthy mice during a forelimb pulling task [Data set]. Zenodo. http://doi.org/10.5281/zenodo.3546068 Processed data and source code that can be used to reproduce the neurophysiological recordings experiment can be found in the following repository: https://gitlab.com/lore.ucci/closed-loop-mouse-stroke-simulation The BNM model and all the relevant information to reproduce the simulated BNM data can be found in the following repository:. Data sharing license: This work is shared under a Creative Commons Attribution CC BY 4.0 license (https://creativecommons.org)

